# TOR inhibition drives accumulation of amino acids through transcriptional activation in algae

**DOI:** 10.1101/2025.09.29.679311

**Authors:** Shivani Upadhyaya, Suzanne M. Kosina, Kim Hixson, Athena Schepmoes, Ritesh Mewalal, Yu Zhang, Ronan O’Malley, Leo A. Baumgart, Ian K. Blaby, Mary S. Lipton, Trent R. Northen, Krishna K. Niyogi, Melissa S. Roth

## Abstract

Cellular homeostasis is maintained by the balance between energy production and breakdown and is fundamental to all forms of life. The conserved, ancient target of rapamycin (TOR) kinase is a central metabolic regulator in eukaryotes that integrates carbon and nitrogen to maintain homeostasis and promote growth and development through protein synthesis. While TOR regulatory mechanisms of amino acid accumulation are well known in yeast and mammals, they remain unknown in photosynthetic organisms. Here, we developed the unicellular green alga *Chromochloris zofingiensis* as a simpler model system for understanding TOR function. Multiomics experiments showed that TOR inhibition leads to an increase in amino acid levels independent of hexokinase-mediated glucose signaling. We observed upregulation of selective amino acid biosynthesis pathways at the transcript and protein levels as potential mechanisms driving the increase in amino acids. Transcriptomics and proteomics experiments identified a basic helix-loop-helix (bHLH) transcription factor with rapid upregulation during TOR inhibition. DAP-seq analysis demonstrated that bHLH can bind directly to the promoters of amino acid biosynthesis genes, potentially regulating their transcription in response to TOR inhibition. We found high conservation of the bHLH-binding motif in the genomes of other green algae and plants, suggesting a conserved regulatory mechanism for amino acid biosynthesis across Viridiplantae. Phosphoproteomics experiments also revealed novel conserved targets that are not currently recognized as part of the TOR pathway. Altogether, our findings elucidate the transcriptional regulation of amino acid metabolism and explain how TOR regulates nitrogen metabolism to support growth and development in photosynthetic organisms.

**Significance Statement:** Carbon and nitrogen metabolism play key roles in enhancing plant yield and reducing fertilizer use. Thus, improving nitrogen utilization can significantly boost crop productivity and algal biotechnology. From yeast to plants to mammals, the protein target of rapamycin (TOR) kinase is an essential metabolic regulator. Here, we developed the unicellular green alga *Chromochloris zofingiensis* as a simpler system to study conserved mechanisms in TOR signaling. Using a multiomics approach, we showed transcriptional regulation of amino acid accumulation upon TOR inhibition and identified a transcription factor with evolutionarily conserved DNA binding sites in nitrogen metabolism genes. We also discovered novel conserved targets of TOR. Our study demonstrates the role of TOR in regulating nitrogen metabolism to support growth and development in photosynthetic organisms.

## Introduction

Energy and nutrient signaling pathways are fundamental regulatory networks controlling growth and development. Integrating multiple nutrient and energy sources, particularly carbon and nitrogen, is essential for maintaining cellular homeostasis. Target of rapamycin (TOR) kinase is a central, conserved metabolic regulator that plays key roles in coordinating carbon and nitrogen sensing and metabolism with signaling pathways that drive growth and development in eukaryotes (1–3). Dysfunctional TOR kinase leads to disease, and nonfunctional TOR kinase is lethal (4). TOR kinase functions as two main complexes, TORC1 and TORC2, in eukaryotes. Both TOR complexes are found in yeast and mammals, whereas photosynthetic organisms only have TORC1. TORC1 is comprised of the TOR kinase, RAPTOR (regulatory associated protein of TOR), and LST8 (Lethal with Sec Thirteen 8) (4–7). In yeast, plants, and mammals, TORC1 senses carbon as sugars and nitrogen as amino acids to promote growth, and TOR inhibition suppresses growth and causes amino acid accumulation (1, 8, 9).

Tightly regulated TOR activity is critical, because prolonged TOR inactivation is associated with immunosuppression and metabolic dysregulation in mammals (4). In mammals and yeast, the mechanism of TOR inhibition-mediated amino acid accumulation involves TOR inhibition or amino acid starvation-induced general control nonderepressible 2 (Gcn2)-mediated translational activation of the general control nonderepressible 4/Activation Transcription Factor 4 (Gcn4/Atf4), a transcription factor (TF) that upregulates expression of amino acid biosynthesis genes (10). This feedback loop ensures reactivation of TOR kinase once the inhibition or nutrient deprivation ends (4). While photosynthetic organisms have homologs of Gcn2, the downstream Gcn4 TF is missing. Consequently, the mechanism regulating TOR reactivation in photosynthetic organisms remains unknown.

Studying TOR signaling in plants is challenging, because they use light, organic carbon (e.g. glucose, sucrose) and inorganic nitrogen (e.g. NO_3_^-^, NH_4_^+^) sources, each capable of activating TOR (11–14). In the model plant *Arabidopsis thaliana*, the complexity is further highlighted by the different activation mechanisms of TOR in shoot and root meristems (13, 15). Layers of tissue-specific regulation have prevented complete characterization of the upstream and downstream components of TOR signaling in photosynthetic organisms.

As green algae are at the evolutionary base of Viridiplantae, they offer simpler single-cell systems for studying TOR signaling in photosynthetic organisms. The TORC1 components are conserved in algae including the model green alga *Chlamydomonas reinhardtii* (Chlorophyceae) (16, 17). Recent studies in *C. reinhardtii* have documented that both photosynthetic CO_2_ fixation and TOR inhibition lead to amino acid accumulation (18, 19). In *C. reinhardtii*, TOR inhibition triggers de novo amino acid synthesis by increasing the activities of glutamine synthetase (GS) and glutamate synthase (19), whereas in *A. thaliana* and the red alga *Cyanidioschyzon mereolae*, TOR inhibition leads to upregulation in gene expression of GS, ammonium (AMT) and nitrate transporters (NRT), and nitrate reductase (NR), thereby suggesting a transcriptional regulation of nitrogen metabolism (20, 21). Because *C. reinhardtii* cannot utilize exogenous glucose, it has limited ability to distinguish TOR-dependent carbon and nitrogen signalling.

The unicellular green alga *Chromochloris zofingiensis* (Chlorophyceae) is an emerging model organism for photosynthesis, metabolism, and energy sensing and signaling (22–28). *C. zofingiensis* readily metabolizes exogenous glucose, a known activator of TOR. When *C. zofingiensis* consumes exogenous glucose under iron-limiting conditions, it switches off photosynthesis and accumulates triacylglycerols (TAGs) via hexokinase1 (HXK1) (23–25). Nitrogen deprivation also suppresses photosynthesis while TAGs accumulate in *C. zofingiensis* (26). Because *C. zofingiensis* accumulates biofuel precursors such as TAGs and the high-value antioxidant astaxanthin while increasing biomass, this alga and its biology are relevant for commercialization (23, 25, 27, 28). In this work, we present *C. zofingiensis* as a single-cell system for elucidating fundamental mechanisms related to TOR signaling. Here, we show that inhibition of TOR increases amino acid biosynthesis through transcriptional activation. Furthermore, we reveal a key TF with conserved DNA binding sites in nitrogen and starch metabolism genes and novel conserved targets of the TOR pathway in photosynthetic organisms.

## Results and Discussion

### TOR Inhibition Prevents Glucose-driven Growth

The TOR signaling pathway regulates growth in response to environmental cues in eukaryotes (4), and TOR inhibition is a critical strategy to determine key players and elucidate the pathway (11-13). We used the kinase inhibitor AZD8055 (AZD), the common ATP-competitive active site TOR inhibitor (7), to inhibit TOR in *C. zofingiensis*. We first determined that 1 µM AZD was the appropriate concentration for *C. zofingiensis*, as it prevented growth on plates (*SI Appendix*, Fig. S1A). Consistent with the literature, this AZD concentration is the same as is used with *C. reinhardtii* (20, 29–31).

To disentangle glucose-TOR signaling in *C. zofingiensis*, we conducted an experiment with TOR inhibition by 1 µM AZD, 100 mM glucose addition, and providing both 100 mM glucose + 1 µM AZD (glucose+AZD) to wild-type (WT) photoautotrophic *C. zofingiensis* cells under continuous illumination (100 µmol photons m^-2^ s^-1^). Cells in all three treatments turned orange while controls remained green (Fig. 1A). As previously reported with glucose, *C. zofingiensis* cells turn orange due to the accumulation of ketocarotenoids (23-25, 27, 28, 32, 33).

**Fig. 1.**
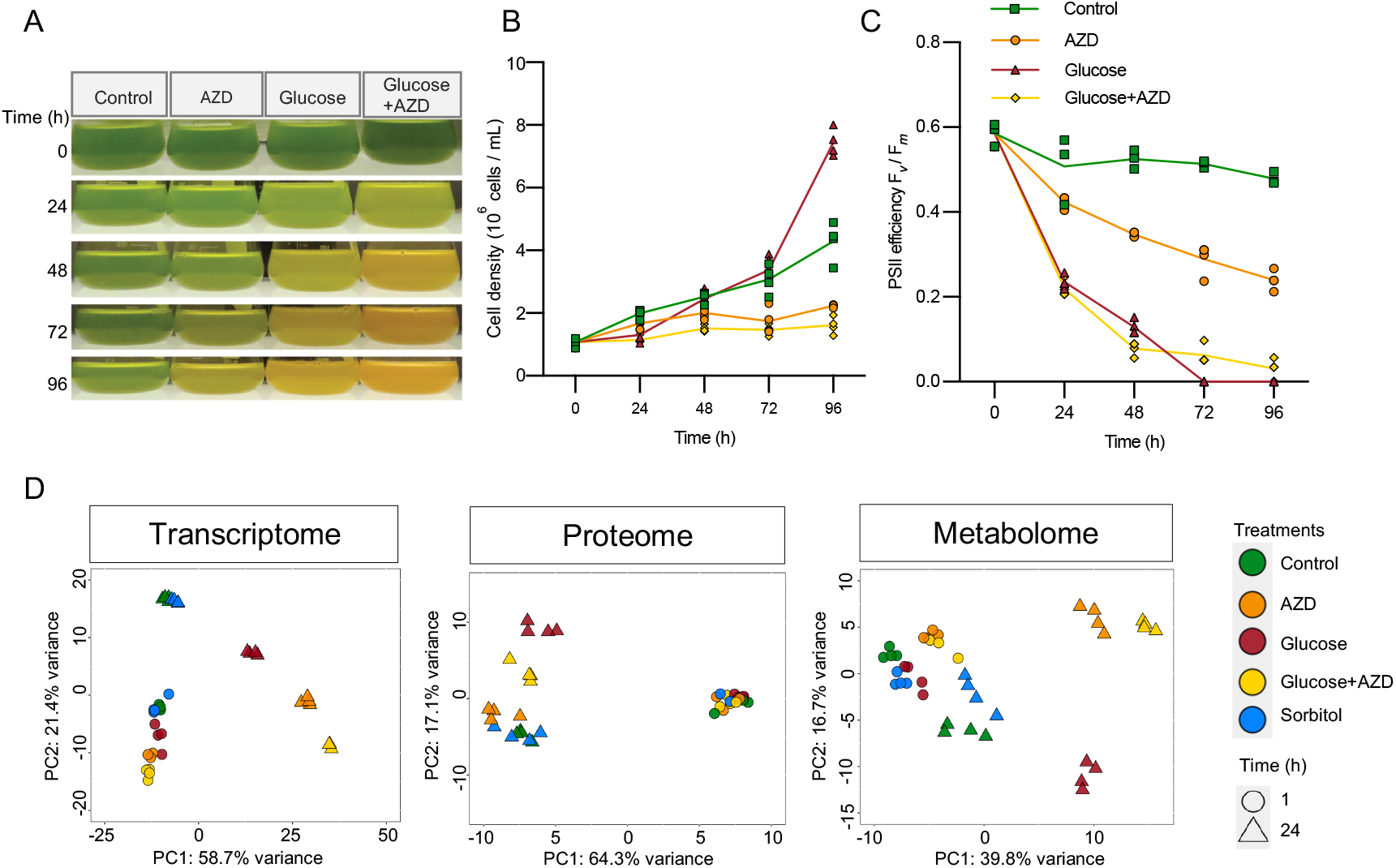
TOR inhibition prevents glucose-driven growth and suppresses photosynthesis in the green alga *C. zofingiensis*. (A) Representative *C. zofingiensis* (WT) cultures over the 96 h experiment with the following treatments: control, TOR inhibition by 1 µM AZD8055, 100 mM glucose addition, and 100mM glucose + 1 µM AZD treatment. (B) Cell density and (C) maximum PSII photosynthetic efficiency (F_*v*_/F_*m*_). For (B) and (C), datapoints represent biological replicates with lines representing means (*n =* 3). Culture biomass shown in Fig. S1B. (D) PCA of WT 1 h and 24 h transcriptome, proteome, and metabolome with treatments: control, 1 µM AZD, 15 mM glucose, 15 mM glucose + 1 µM AZD, and 15 mM sorbitol as an osmotic control. For the transcriptome, normalized log_2_-transformed counts of 500 genes with highest variance were considered. For the proteome and the metabolome, the raw intensity values were log_2_-transformed, and proteins or metabolites with highest variance were used to calculate and plot the PCA. Data represent four biological replicates.

TOR inhibition, independently and in combination with glucose, led to decreased photosynthesis and prevented glucose-dependent increases in cell density and culture volumetric biomass (Fig. 1B-C, *SI Appendix*, Fig. S1B), showing that TOR inhibition prevents growth and suppresses photosynthesis in *C. zofingiensis*. These results are consistent with previous studies in *C. reinhardtii* and *A. thaliana* (30, 34, 35). Glucose addition alone switched off photosynthesis and caused an increase in cell growth and biomass (Fig. 1A-C, *SI Appendix*, Fig. S1B), confirming that glucose is a significant driver of fast growth and high biomass accumulation in *C. zofingiensis*, as shown previously (23-25, 28). However, our results demonstrated that TOR inhibition blocks glucose-driven biomass accumulation, indicating that TOR inhibition overrides or blocks glucose signaling.

### Distinct Regulation of TOR Inhibition and Glucose Responses

To decipher the independent and synergistic signaling of TOR inhibition and glucose in *C. zofingiensis*, we performed transcriptomics, TMT proteomics, and metabolomics experiments with the following treatments: TOR inhibition by 1 µM AZD, 15 mM glucose addition, the combined effect of 15 mM glucose + 1 µM AZD, and 15 mM sorbitol addition (as an osmotic control) at 1 h and 24 h time points (*SI Appendix*, Fig. S1C).

Principal component analyses (PCA) of the omics datasets showed that TOR inhibition and glucose addition, both independently and synergistically, significantly altered the transcriptome, proteome, and metabolome particularly by 24 h (Fig. 1D). While the transcriptome PC1 accounted for 58.7% of the variance and was driven by glucose+AZD at 24 h, the proteome PC2 accounted for 17% of the variance and was driven by glucose at 24 h. The AZD and glucose+AZD proteomes clustered near the control, indicating that a large portion of the proteome did not change significantly despite the transcriptome showing significant variance for AZD treatments. These results suggest significant modulation of the transcriptome that is not reflected in the proteome. TOR is also a known regulator of global translation in algae, plants, yeast, and mammals and regulates the translation machinery via phosphorylation at multiple levels (36), and thus, TOR inhibition can lead to a decrease in overall translation and dampening of the proteome. Furthermore, the metabolome PCA showed that while AZD and glucose+AZD metabolites clustered together, PC1 and PC2 only explained ∼45 % of the variance.

Overall, our results indicate that AZD and glucose have distinct transcriptional and metabolic responses, while glucose+AZD modulates the transcriptome synergistically. Given the transcriptomic and metabolomic changes observed during TOR inhibition and glucose addition, we next explored the downstream signaling by TOR kinase, through phosphoproteomic profiling.

### Identification of Novel TOR Players

To confirm AZD-mediated inhibition of TOR and to identify known and uncharacterized TOR targets in *C. zofingiensis*, we performed phosphoproteomics of TOR inhibition using the same treatments described above (1 µM AZD, 15 mM glucose addition, 15 mM glucose + 1 µM AZD, and 15 mM sorbitol-treated WT cultures at 1 h and 24 h). Because glucose has previously been shown as an upstream activator of TOR in plants (11), we hypothesized that TOR-dependent sites of protein phosphorylation would be downregulated during TOR inhibition in both AZD and glucose+AZD treatments, while glucose would upregulate the same phosphosites. Proteins were extracted and phosphopeptides were further enriched using TiO_2_ magnetic beads for detection by Tandem Mass Tag Mass Spectrometry (TMT MS). This experiment led to the detection of 19,030 phosphopeptides on 3,861 proteins with a peptide being detected in at least one treatment (Dataset S1). To extract statistically significant and TOR-dependent phosphopeptides, we performed hierarchical clustering on selected peptides whose log_2_ fold changes (log_2_FC) in AZD were ≥ ±1 and with a *P* ≤ 0.01 (Student’s *t*-test).

With this approach, we identified 1288 significant phosphopeptides. To uncover the underlying biological processes in each cluster, we conducted *k*-means clustering with four centers and gene ontology (GO) enrichment analysis (Fig. 2A, *SI Appendix*, Fig. S2A, Dataset S2-S3). Cluster 1 showed upregulated phosphopeptides at both 1 h and 24 h in all treatment conditions and was enriched in terms associated with phosphoinositol biosynthesis and ADP-ribosylation factor (ARF) signal transduction. Cluster 2 showed downregulation at 1 h in all treatments and was enriched in terms associated with photosynthesis and cytoskeleton organization. Cluster 3 showed upregulation at 1 h and downregulation at 24 h in all treatments and was enriched in terms associated with amino acid catabolism and carbohydrate catabolism. Distinctively, cluster 4 showed downregulation during TOR inhibition in AZD at 1 h and was enriched in terms associated with TOR signaling, tRNA modification, and chromatin remodeling, which suggests that cluster 4 includes peptides specifically regulated by TOR (Fig. 2, *SI Appendix*, Fig. S2A, Dataset S3).

**Fig. 2.**
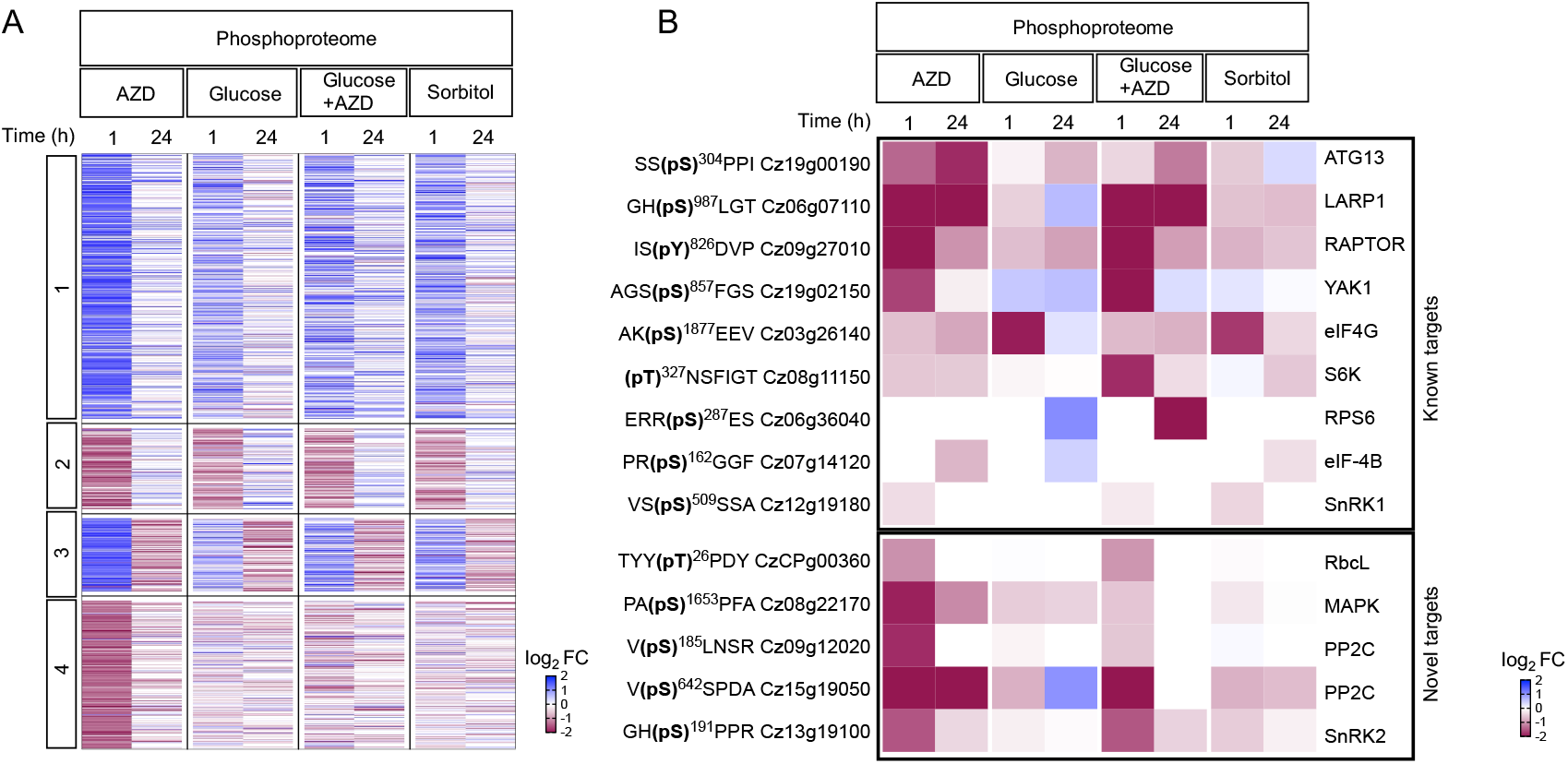
Phosphoproteomics of TOR-inhibited *C. zofingiensis* identifies novel TOR players. (A) The log_2_ fold changes of phosphopeptides between TOR inhibition by 1 µM AZD, 15 mM glucose addition, 15 mM glucose + 1 µM AZD, and 15 mM sorbitol-treated *C. zofingiensis* (WT) cultures at 1 h and 24 h were clustered into four groups using *k*-means clustering. (B) The log_2_ fold changes of TOR-dependent phosphosites from cluster 4 were labelled according to their protein symbols and phosphorylated Ser/Thr residue on the left and by their ortholog name identified on the right. The phosphopeptides were labelled if they are known conserved TOR targets as “known targets” and if they are unknown as “novel targets”. RbcL, MAPK, PP2C, PP2C, and SnRK2 were identified as putative novel conserved TOR targets. White boxes in the heatmap indicate that the phosphopeptides were not detected. Data represent means of fold changes log_2_ intensities of phosphopeptides (*n* = 3-4 biological replicates).

To verify phosphopeptides affected by TOR inhibition, we identified orthologs of several known TOR-dependent proteins found in the literature from plants and mammals. These included conserved and experimentally characterized targets of TOR phosphorylation such as the autophagy initiating complex protein 13 (ATG13), the La-related protein 1 (LARP1), the ribosomal protein S6 kinase (S6K), and protein translation-associated eukaryotic initiation factors (eIF4G and eIF4B) (12, 35, 37, 36). Phosphopeptides of ATG13, LARP1, S6K, eIF4G, and eIF4B were found in cluster 4 and downregulated in both AZD and glucose+AZD treated cultures while LARP1 and eIF4B were upregulated in glucose addition by 24 h (Fig. 2B). These findings highlight the conserved role of TOR in regulating protein biosynthesis, similar to what has been observed in plants (36). The downregulation of these phosphopeptides in both glucose- and sorbitol-treated cultures at 1 h may suggest an effect of osmotic shock on these phosphopeptides.

Cluster 4 also included phosphopeptides from the substrate of S6K (ribosomal protein S6 (RPS6)), Yet Another Kinase 1 (YAK1), RAPTOR, and Snf1-related kinase 1 (SnRK1) (Fig. 2B). In *A. thaliana*, S6K, RPS6, and YAK1 are phosphorylated by TOR kinase (34, 38–40), while RAPTOR and SnRK1 are known to interact with the TOR complex to regulate energy sensing and signaling (12). Our proteomics dataset showed that there were no changes in the total protein abundances of the conserved targets, which suggests that TOR regulates a specific post-translational response in *C. zofingiensis* (*SI Appendix*, Fig. S2B). In total, cluster 4 of our phosphoproteome analysis revealed 206 differentially expressed phosphopeptides from 164 proteins, 87% of which have a GO term and 13% with no assigned GO term (*SI Appendix*, Dataset S2).

Intriguingly, we discovered conserved proteins not previously associated with TOR signaling in the TOR-specific cluster 4 (Fig. 2B, Dataset S5). This experiment revealed putative novel TOR targets including Rubisco large subunit (RbcL), mitogen-activated protein kinase (MAPK), two protein phosphatase 2C (PP2C), and SnRK2. Although these proteins are well studied, their connections to TOR signaling have not been previously reported. Furthermore, we identified phosphosites on these proteins that were dephosphorylated within 1 h of AZD and glucose+AZD. PP2C (Cz15g19050) also showed an upregulation in phosphorylation with glucose at 24 h (Fig. 2B), suggesting its phosphorylation by glucose-TOR signaling.

It is curious that we identified RbcL as a potential target of TOR. In *C. reinhardtii*, the diphosphoinositol pentakisphosphate kinase (VIP1) uses inositol pyrophosphates (InsP_7_/InsP_8_) to regulate phosphorylation of photosystem II core proteins via the VIP1-TOR axis (51). This pathway also controls phosphorylation of the kinase state transition like (Stl1), a homolog of the *A. thaliana* state transition kinase 8 (STN8). Since STN8 phosphorylates chloroplast proteins like RbcL (52), VIP1-TOR could indirectly regulate RbcL phosphorylation via Stl1. However, this proposed TOR–VIP–Stl1/STN8 axis regulating RbcL requires further validation. In *A. thaliana*, the TOR complex has also previously been reported to associate with proteins involved in chloroplast development, as well as the chloroplast-targeted Chaperonin 60 (Cpn60), suggesting a role for TOR in regulating chloroplast function (12, 34, 35). However, the underlying mechanisms governing this interaction remain to be elucidated. While further experimental validation is needed to establish these proteins as downstream direct or indirect targets of TOR, these results suggest roles for RbcL, MAPK, PP2C, and SnRK2 in the TOR pathway in

*C. zofingiensis*.

Overall, our phosphoproteomic analysis shows a decrease in phosphorylation of conserved TOR targets with AZD treatment, thereby confirming inhibition of TOR in *C. zofingiensis* in our AZD and glucose+AZD treatments. The increase in phosphorylation at the same site during glucose addition also establishes glucose as an upstream activator of TOR in *C. zofingiensis*. In addition to known targets, our phosphoproteomic experiment reveals a chloroplast-encoded protein RbcL and other proteins including MAPK, PP2C, and SnRK2 as novel players in the TOR pathway.

### TOR Inhibition Triggers Upregulation of Amino Acid Biosynthesis Pathways and Accumulation of Amino Acids

To distinguish TOR-dependent and independent responses to glucose, we integrated transcriptomic (1 h) and proteomic (24 h) data for 9,528 transcripts and proteins. Further, *k*-means clustering of differentially expressed transcripts (*P* ≤ 0.01 from DESeq2 analysis) identified 6 clusters that were enriched with GO terms (*SI Appendix*, Fig. S3, Dataset S4, Dataset S5). Cluster 1 of the transcriptome showed significantly upregulated genes in AZD, glucose addition, and glucose+AZD and was enriched in terms associated with glycolysis and lipid metabolism related terms. Clusters 2 and 3 showed downregulated genes in AZD, glucose, and glucose+AZD treatments and were enriched in terms associated with chlorophyll catabolism, cell cycle, and sugar metabolism. Most strikingly, we identified Cluaster 4 as a TOR-specific cluster and transcripts in cluster 4 were upregulated by glucose (indicating TOR activation) and downregulated by either AZD or glucose+AZD (indicating TOR inhibition). This cluster was enriched for translation, rRNA/tRNA processing, and nitrogen metabolism. Cluster 5, enriched for protein catabolism and autophagy, showed upregulation under AZD and glucose+AZD, consistent with stress responses. Lastly, cluster 6 specifically comprised genes that were downregulated solely under glucose conditions and were enriched in terms associated with photosynthesis and pigment biosynthesis and have been characterized previously in *C. zofingiensis* (23, 25). Overall, transcriptomics identified genes that are specifically regulated by TOR and are part of the glucose-TOR signaling pathway in Cluster 4. Additionally, the downregulation of protein biosynthesis and translation-related genes in *C. zofingiensis* during TOR inhibition establishes the role of TOR in translational inhibition as seen in other eukaryotes (36).

Comparing log_2_FC of 9,528 candidate genes with both steady-state transcript (1 h) and protein levels (24 h) across AZD, glucose, and glucose+AZD treated samples (Dataset S6), revealed a key role for TOR in regulating translation. We mapped the relationships into scatter plots that were further divided into four sectors: only upregulated in mRNA (orange), upregulated in both mRNA and protein (blue), only downregulated in mRNA (gray) and downregulated in both mRNA and protein (green) (Fig. 3) and performed pathway enrichment for each sector. The transcript levels showed the highest positive correlation with the protein levels in glucose-treated samples with significant protein increases in fatty acid biosynthesis and glycolytic pathways (blue) and a simultaneous downregulation of proteins in the chlorophyll biosynthetic pathway (green) (R^2^ = 0.47, Fig. 3B, Dataset S7), which is consistent with previous studies in *C. zofingiensis* (23, 25). Transcripts involved in pathways such as sucrose biosynthesis, tRNA charging, and starch synthesis were upregulated (orange). We observed a decreased correlation for mRNA and protein in AZD (R^2^ = 0.22) and glucose+AZD (R^2^ = 0.38), suggesting an inhibition of translation during TOR inhibition independent of glucose (Fig. 3). Surprisingly, mRNA and proteins of both glycolysis and fatty acid biosynthetic pathways increased in glucose+AZD, indicating that transcription was TOR independent. Although we observed an overall dampening of translation during TOR inhibition in AZD treatment conditions, glutamine biosynthesis and nucleotide degradation pathways were upregulated at the protein level (blue, Fig. 3). Additionally, we detected valine, isoleucine and threonine biosynthetic pathways were also upregulated (orange) at the mRNA level in both AZD and glucose+AZD (Fig. 3, Dataset S7). Altogether, TOR inhibition reduced overall translation even in the presence of glucose, while concomitantly upregulating amino acid biosynthetic pathways at the mRNA level.

**Fig. 3.**
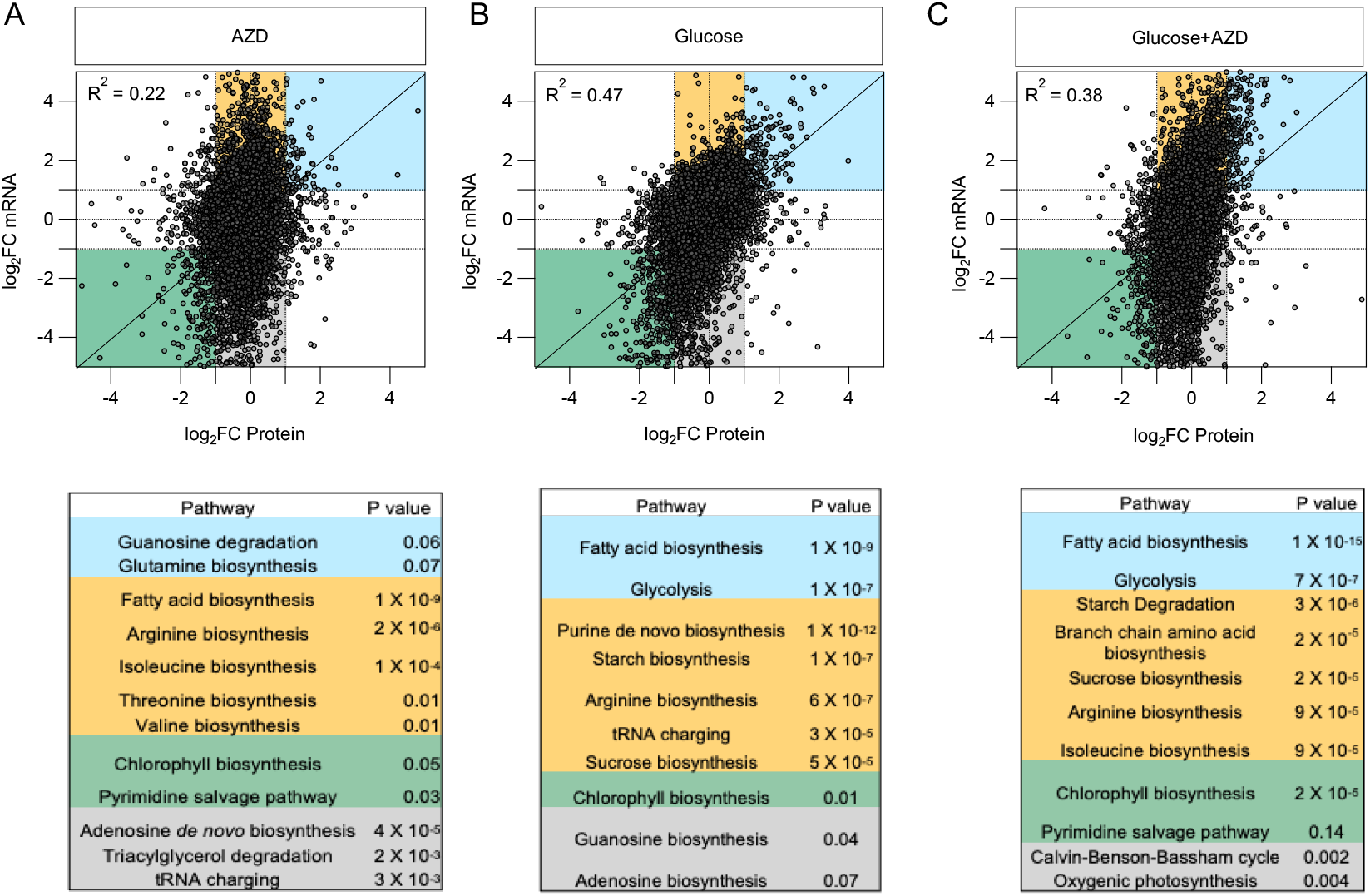
Integrating transcriptomics and proteomics reveals TOR inhibition-dependent upregulation of amino acid pathways. Scatterplot representation of correlation of mRNA and protein expression ratios. The log_2_ value of *C. zofingiensis* (WT) mRNA at 1 h and protein at 24 h ratios of expression during (A) TOR inhibition by 1 µM AZD, (B) 15 mM glucose addition, and (C) 15 mM glucose + 1 µM AZD treatment conditions. The scatterplots were divided into four sectors with upregulated mRNA (orange), upregulated in both mRNA and protein (blue), downregulated mRNA (gray), and downregulated in both mRNA and protein (green). The coefficient of correlation was plotted as R^2^. Pathway enrichment was conducted as described in *SI Appendix*, Materials and Methods and Dataset S7 for the complete list. Data represent means of fold changes of log_2_ counts of transcripts and log_2_ intensities of proteins (*n* = 3-4 biological replicates).

To determine if the TOR inhibition dependent upregulation of amino acid biosynthesis pathways led to amino acid accumulation, we clustered our untargeted data and performed *k*-means clustering (five clusters) on the putatively annotated features (S*I Appendix*, Fig. S4, Dataset S8). Remarkably, we observed an increase in amino acids solely during TOR inhibition in cluster 1, evident as early as 1 h and consistently observed at 24 h. The increase in amino acids during TOR inhibition is consistent with previous studies in *C. reinhardtii* and *A. thaliana* (19, 21). Clusters 2 and 3 of the metabolome showed an accumulation of metabolites such as lipids and nucleobases in AZD, glucose, and glucose+AZD treatments by 1 h and went down by 24 h. These results were consistent with the transcriptome response, which showed an upregulation in lipid metabolism genes with these treatments (Fig. 3, *SI Appendix*, cluster 2, Fig. S3). Cluster 4 showed an accumulation of sugars such as sucrose, trehalose, lactulose, and more in glucose-treated cultures independent of TOR inhibition. Cluster 5 represented a treatment non-specific reduction of metabolites including glycerolipids. To elucidate how amino acids are highly upregulated during TOR inhibition by 1 h, we further explored the amino acid biosynthesis pathways.

De novo amino acid biosynthesis relies on precursors derived from the central metabolic pathways of glycolysis and the tricarboxylic acid (TCA) cycle (Fig. 4A). While the pathways listed in Fig. 4A are not comprehensive, key precursors and fueling reactions for de novo amino acid biosynthesis include pyruvate for biosynthesis of alanine, leucine and valine; α-ketoglutarate for the synthesis of glutamate, proline, arginine and glutamine; and oxaloacetate for the synthesis of aspartate and related amino acids (41, 42). Phosphoenolpyruvate, a glycolytic intermediate, acts as a precursor for aromatic amino acids, while the pentose phosphate pathway intermediates, ribose 5-phosphate and erythrose 4-phosphate, act as precursors for the synthesis of histidine and aromatic amino acids, respectively. Inorganic nitrogen is also assimilated into glutamate- and glutamine-derived amino acids (42).

**Fig. 4.**
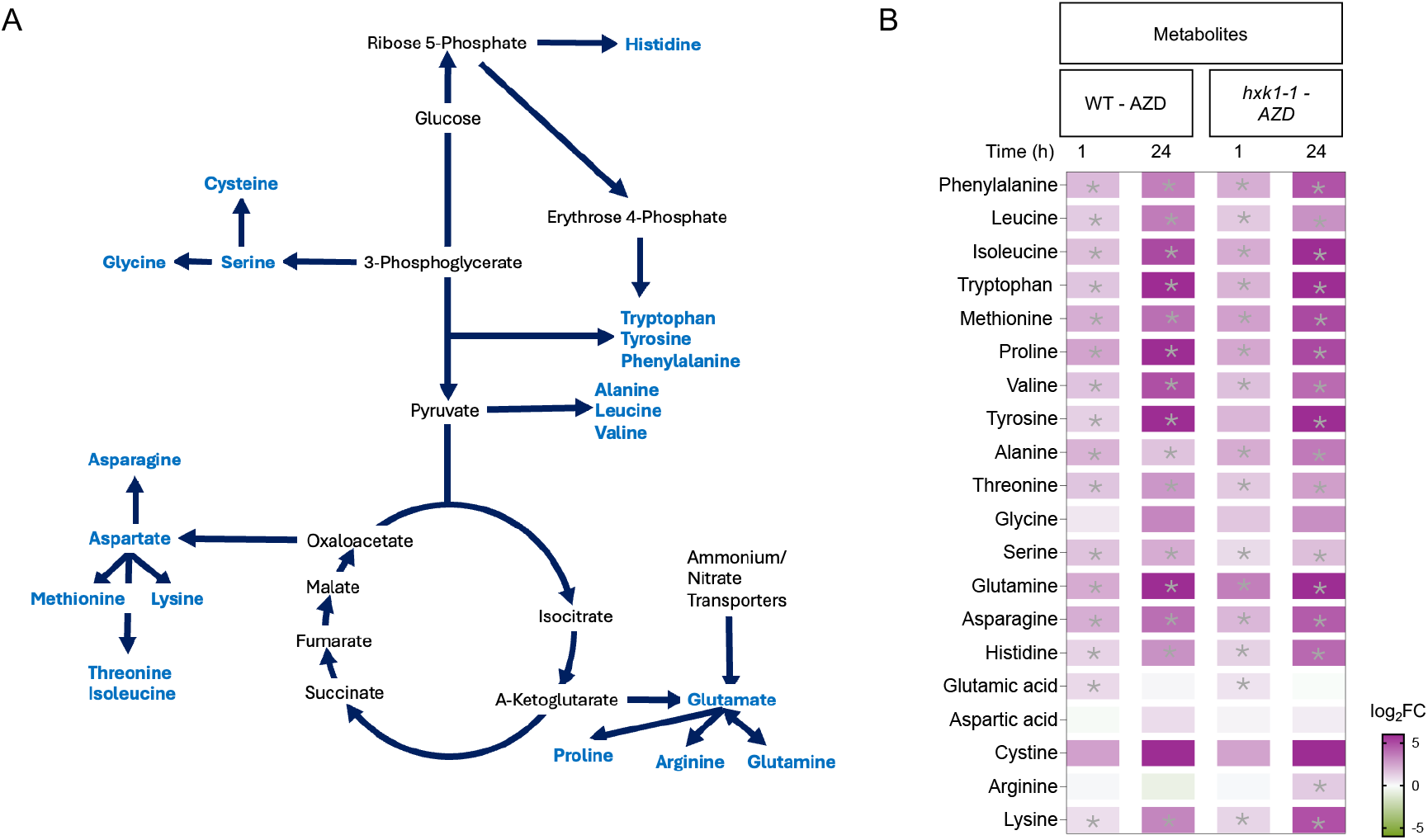
TOR inhibition increases amino acids independent of hexokinase1 (HXK1). (A) Representation of amino acid biosynthesis precursors and pathways. Light blue represents amino acids and black represents the carbon skeletons used for their biosynthesis. (B) Log_2_ fold changes (FC) of amino acids in *C. zofingiensis* WT and *hxk1-1* mutant during TOR inhibition by 1 µM AZD at 1 h and 24 h. Data represent mean fold changes of log_2_ intensities of metabolites (*n* = 3-4 biological replicates) and asterix (*) represents *P* ≤ 0.01 (Student’s *t*-test).

Targeted metabolite analysis of TOR-inhibited WT cells revealed a significant increase in many amino acids at both 1 h and 24 h (*P* ≤ 0.01, Student’s *t*-test) (Fig. 4B). We further analyzed the transcript and protein log_2_ fold changes of the above-mentioned biosynthetic pathways in WT during TOR inhibition by 1 µM AZD, 15 mM glucose addition, and 15 mM glucose + 1 µM AZD treatment (*SI Appendix*, Fig. S5, Dataset S9). We observed a strong upregulation of the amino acid biosynthetic pathways at the transcript level but selective upregulation at the protein levels. Specifically, enzymes involved in the biosynthesis of aspartate, asparagine, glutamate and glutamate derived, and TCA cycle derived were upregulated at the transcript and protein levels in AZD and glucose+AZD by 24 h. However, the enzymes associated with branched chain amino acids (leucine and valine) were upregulated at the transcript but not at the protein level in AZD treatment by 24 h. However, leucine and valine accumulated in TOR-inhibited samples (Fig. 4B), suggesting that the translation of their biosynthetic enzymes is sufficient to promote accumulation or the involvement of either translation inhibition or protein catabolism, possibly via the proteasomal pathway. To explore alternative mechanisms of amino acid accumulation, we conducted a 1 h experiment of TOR inhibition by 1 µM AZD, 35 µM cycloheximide (CHX) treatment to inhibit cytoplasmic protein synthesis, 10 µM MG-132 treatment to prevent proteasome-dependent protein degradation, and a combination of TOR inhibition + 35 µM CHX or 10 µM MG-132 following a strategy previously used in *C. reinhardtii* (19). Consistent with previous studies, we observe that both translation inhibition (CHX) and TOR inhibition (AZD) cause substantial increases in log_2_FC of free amino acid levels, with combined treatment leading to a further elevation (*SI Appendix*, Fig. S7, Dataset S11). However, in our treatment, TOR inhibition specifically increases log_2_FC of alanine, glutamine, glutamic acid, arginine, glycine, asparagine, histidine, lysine, and serine levels (*P* ≤ 0.01, Dunnett’s test) in comparison to only CHX treatment, suggesting their accumulation is by de novo biosynthesis instead of protein turnover. Proteasome inhibition with MG-132 alone had limited impact on the log_2_FC of amino acid levels. However, its combination with AZD resulted in an increase (*P* ≤ 0.01, Dunnett’s test), equivalent to AZD treatment alone (Fig. S7, Dataset S11). Overall, our results indicate that TOR inhibition based amino acid accumulation occurs through non-proteasomal proteolytic pathways, in addition to the upregulation of de novo biosynthesis. To untangle TOR signaling from glucose-HXK signaling we used the *C. zofingiensis* HXK1 mutant (*hxk1-1*). This mutant was generated by forward genetics, has a strong HXK1 knockdown phenotype, and during growth on glucose it maintains photosynthesis and does not accumulate astaxanthin, TAGs, or cytoplasmic lipid droplets (24, 25). These studies establish that the increase in biomass and inhibition of photosynthesis is glucose-and HXK1-dependent (24, 25). Metabolite analysis showed that TOR inhibition (AZD) also increased amino acid levels in the *hxk1-1* mutant (Fig. 4B), demonstrating that amino acid accumulation is a direct consequence of TOR inhibition and independent of the glucose-HXK1 signaling axis. Furthermore, transcript and protein analysis revealed that while TOR inhibition upregulated amino acid biosynthetic pathways in *hxk1-1* similar to WT, the glucose-dependent upregulation observed in WT at 24 h was absent in the mutant (SI Appendix, Fig. S6, Dataset S10). These results indicate that glucose stimulation of amino acid biosynthesis requires HXK1. Altogether, our metabolomics data confirmed TOR inhibition-specific accumulation of amino acids is regulated at both the transcript and protein levels, independent of the glucose-HXK axis.

### G2-like, HSF, and bHLH Transcription Factors Bind to Amino Acid Biosynthesis Genes

To identify the underlying transcription factors (TFs) involved in this response in *C. zofingiensis*, we examined differentially expressed TFs that were upregulated under TOR inhibition in transcriptomic clusters 1 and 5 (*SI Appendix*, Fig. S3). We extracted 48 TFs upregulated at either 1 h or 24 h of AZD or glucose+AZD treatments. We then filtered our list to 10 orthologs of the previously identified TFs in nitrogen-responsive networks in *A. thaliana* (47). We hypothesized that TOR inhibition leads to transcriptional activation of amino acid biosynthesis, along with corresponding increases in their amino acid levels (Fig. 4, Fig. S5 & Fig. S6). We performed DNA affinity purification sequencing (DAP-seq) to map genome-wide TF-DNA interactions and identify target promoters. We further hypothesized that the TFs associated with nitrogen-responsive networks might also regulate transcript changes associated with TOR inhibition. We performed DAP-seq on five of the 10 TFs and obtained high quality data for three TFs (Fig. 5).

**Fig. 5.**
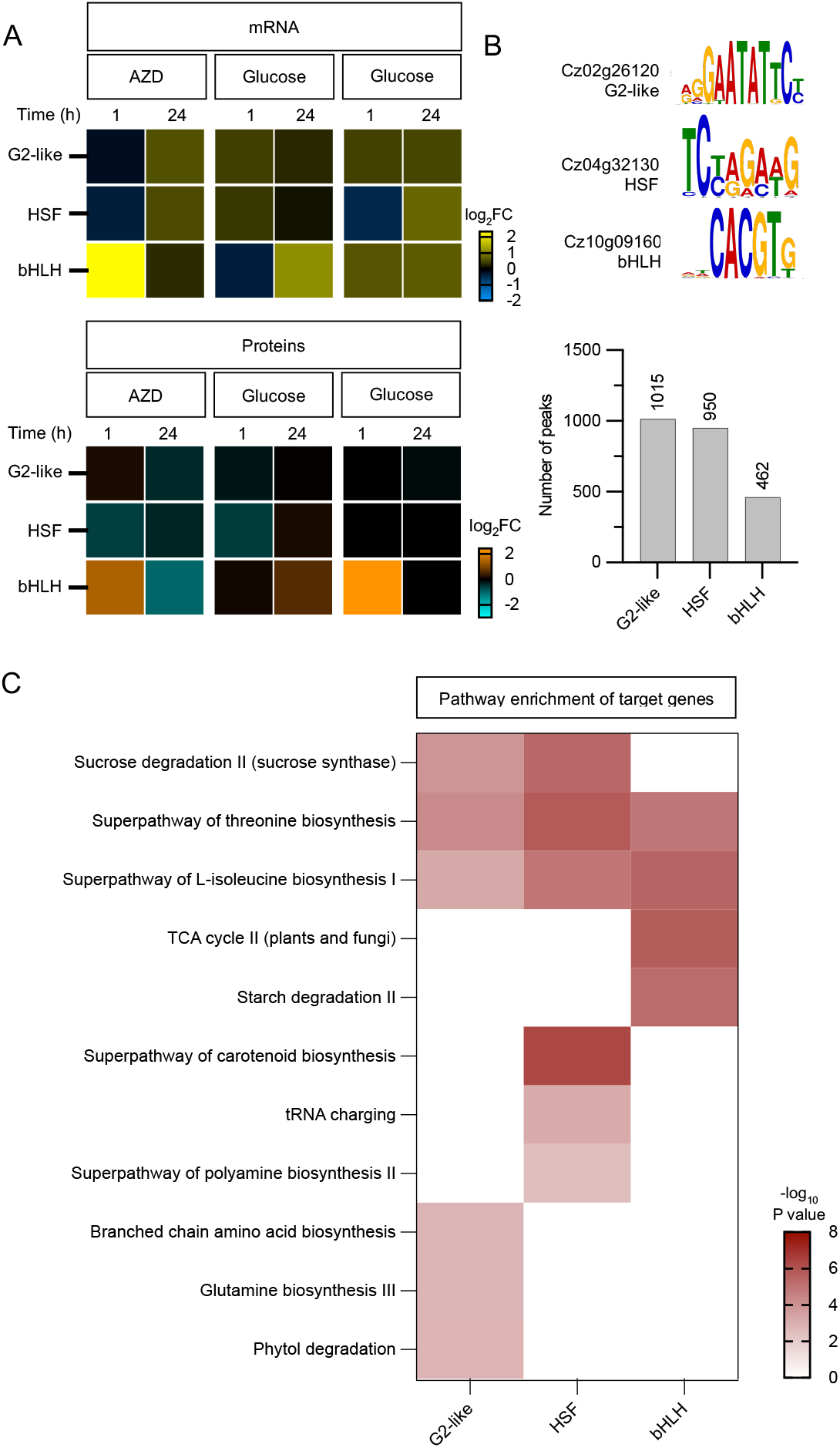
G2-like, HSF, and bHLH transcription factors are associated with amino acid biosynthesis pathways. (A) mRNA and protein log_2_FC of G2-like (Cz02g26120), HSF (Cz04g32130) and bHLH (Cz10g09160) TFs during TOR inhibition by 1 µM AZD. Data represent log_2_FC of mRNA counts (*n* = 3-4 biological replicates). (B) DAP-seq based motif identification for G2-like, HSF, and bHLH TFs and the number of their target peaks. (C) Heatmap displaying the −log_10_ P-value for enriched pathways of the target genes associated with the three TFs.

The three TFs: a Golden2-like (G2-like) domain TF (Cz02g26120), a heat shock factor (HSF, Cz04g32130), and a basic helix loop helix (bHLH) domain-containing TF (Cz10g09160), were upregulated during TOR inhibition at both transcript (*P* ≤ 0.01 from DESeq2 analysis, Dataset S4) and protein levels (Fig. 5A). De novo motif discovery of the most enriched peaks bound by the respective TF found that the most significant motifs were 8-11 nucleotides long (Fig. 5B). The bHLH TF was bound to two adjacent motifs, each approximately 6 bp long. The number of target genes bound by each TF was ∼1000 genes (Fig. 5B, Dataset S13).

To gain insight into the potential biological functions of the three TFs and their target genes, we performed a pathway enrichment analysis for the targets of each (Dataset S14). Like GO enrichment, pathway enrichment can be used to enrich biological pathways associated with the target genes of the selected TFs. Enrichment in the isoleucine and threonine biosynthesis pathways was observed for all three TFs (Fig. 5C). Both the G2-like and the HSF TF were also found to bind to glutamine synthetase, aspartate aminotransferase, nitrate and ammonium transporter genes, which showed upregulation during TOR inhibition in both WT and *hxk1-1* mutant (*SI Appendix*, Fig. S8). TOR kinase-mediated transcriptional regulation is well understood in other eukaryotes. In yeast, YAK1 phosphorylates and regulates transcription factors such as HSF during nutrient starvation (43). In *A. thaliana*, the kinase YAK1 interacts with RAPTOR and regulates nitrogen metabolism (44). In *A. thaliana*, YAK1 inhibition in TOR-deficient mutants reduces glutamine levels and restores growth, suggesting a conserved role for YAK1 in regulating nitrogen metabolism. Therefore, a similar regulation of the TOR-RAPTOR-YAK1-HSF may be involved in regulating nitrogen metabolism in plants and algae.

### bHLH Transcription Factor May Regulate Central Metabolism, Nitrogen, and Starch Metabolism

The bHLH TF was strongly upregulated at both transcript and protein levels by 1 h of TOR inhibition and binds to genes related to the TCA cycle, amino acid biosynthesis, and starch metabolism (Fig. 5). Although DAP-seq data cannot confirm transcriptional activation, the binding patterns suggest regulatory roles. Thus, these results suggest bHLH is a key TOR-regulated factor involved in the transcription of TOR-responsive genes.

It is well known that most TFs regulate transcription by binding specifically at the promoter region of genes (45, 46). To check if the target genes are specifically bound by the bHLH TF at the promoter and are transcriptionally upregulated during TOR inhibition, we examined the genomic binding patterns and the transcript fold change after 1 h of TOR inhibition. Our pathway enrichment data showed that the bHLH target genes fell into two primary categories: 1) those related to nitrogen metabolism (including genes in the TCA cycle, amino acid biosynthesis and various transporters), and 2) those related to starch degradation (Fig. 5C).

DAP-seq data revealed that the bHLH TF binds to the promoter region (±1500 bp from the transcription start site) of nitrogen metabolism and starch metabolism genes (Fig. 6). We observed that bHLH binds to the promoter of nitrogen metabolism genes including the NADH-glutamate synthase, malate dehydrogenase, isocitrate dehydrogenase, succinate dehydrogenase, alanine transaminase, L-aspartate oxidase, threonine synthase, and allophanate hydrolase (Fig. 6B). These genes were also highly upregulated during TOR inhibition (Fig. 6A). We also noticed that the bHLH TF binds to the promoters of starch degradation genes including glucose 1-phosphate adenylyltransferase, and beta amylase, which were also highly upregulated during TOR inhibition. In *A. thaliana*, an ortholog of the bHLH TF was identified as a potential candidate in nitrogen metabolism from a large-scale transcriptional network analysis (47). Moreover, in *A. thaliana* SnRK1 also interacts with bHLH4 TF in response to nitrogen levels, suggesting a role of bHLH TFs in energy metabolism (48, 49).

**Fig. 6.**
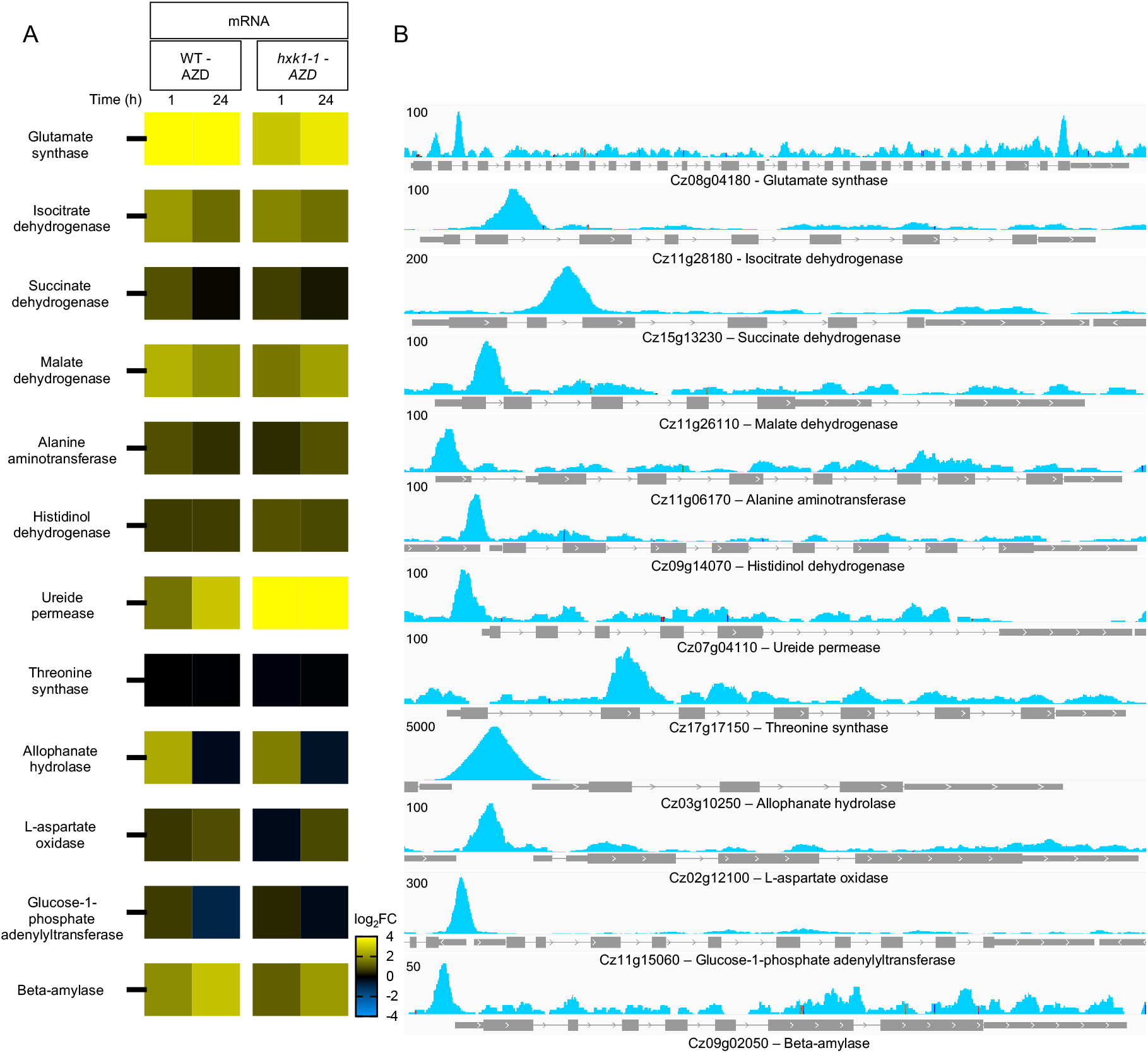
Binding of bHLH to nitrogen and starch metabolism genes potentially regulates transcription. (A) Heatmap of nitrogen and starch metabolism gene expression during TOR inhibition by 1 µM AZD in WT and *hxk1-1* mutant. Data represent mean fold changes (FC) of log_2_ mRNA counts (*n* = 3-4 biological replicates). (B) DAP-seq peaks in blue associated with nitrogen and starch metabolism genes. Peak height number represents the number of reads at the genomic location or the strength of TF binding. Gene architecture with gray boxes indicating exons, lines indicating introns, and gaps indicating intergenic regions.

While this study and previous studies (47-49) suggest that bHLH TFs interact with TOR and/or its associated signaling pathways, further experiments are needed to dissect how. The precise mechanism by which TOR inhibition upregulates bHLH at the transcript and protein levels (Fig. 5A) and leads to increased amino acid levels remains unknown. Crosstalk between glucose-HXK1 and TOR signaling can also upregulate amino acids, perhaps via the bHLH TF, and future studies can address this aspect. Overall, our results indicate that the bHLH TF specifically binds to the promoters of TCA cycle, nitrogen, and starch metabolism genes to upregulate their transcription during TOR inhibition.

### bHLH Transcription Factor Binding Motif is Conserved in Nitrogen and Starch Metabolism Genes in Algae and Plants

The bHLH family TFs recognize the core DNA sequence motif known as the E-box (5’-CANNTG-3’), of which the most common is the palindromic G-box (5’-CACGTG-3’) (50). We found that the bHLH TF binding motif CACGTG in nitrogen and starch metabolism gene promoters (Fig. 5B). To test the conservation of the binding motif of target gene promoters across various green algae and plants, we identified orthologs of targets related to nitrogen and starch metabolism and scanned their promoters (±1500 bp from the transcription start site) for the presence of the canonical E- or G-box in these orthologs (Dataset S15). In addition to *C. zofingiensis*, we included the green algae *C. reinhardtii, Volvox carteri* (Chlorophyceae), and *Auxenochlorella* sp. UTEX 250-A (Trebiouxiophyceae), and plants *A. thaliana* and *Oryza sativa*. Because our sampling included plants and algae, and algae from Chlorophyceae and Trebiouxiophyceae, which diverged >800 million years ago (53), our sampling covers deep evolutionary distance.

Almost all genes (10/12) had an ortholog with the E/G-box within their promoter regions in the sampled algae as well as plants (Fig. 7). Intriguingly, we could not identify orthologs for allophanate hydrolase suggesting that this enzyme might be specific to *C. zofingiensis* and closely related algae. Another exception was the L-aspartate oxidase, which had an ortholog present in *A. thaliana* and the rice *O. sativa*, but an E/G-box was not identified in its promoter region. However, all three TCA cycle genes that are important for amino acid biosynthesis and nitrogen metabolism had ortholog promoters with the conserved motifs. Overall, these data show high conservation of the E/G-boxes that are bound by bHLH transcription factors and could indicate a conserved mechanism of regulation of these genes by bHLH TFs.

**Fig. 7.**
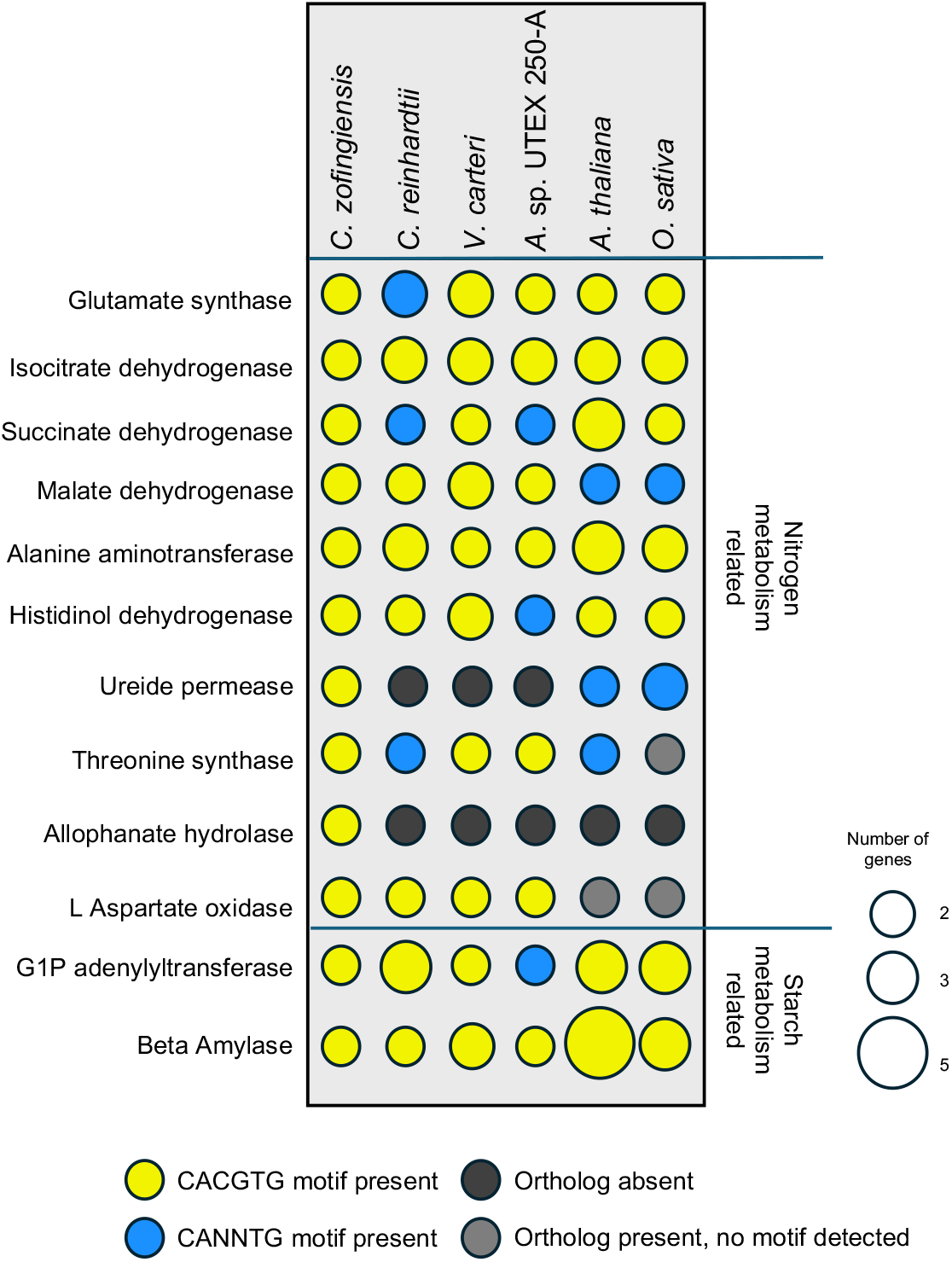
The bHLH binding motif is conserved among photosynthetic organisms. Comparative plot for the presence or absence of the two bHLH DNA binding motifs CACGTG/CANNTG in the promotors of orthologs of nitrogen and starch metabolism genes from Fig. 6.

## Conclusions

We define a pathway regulating the accumulation of amino acids during TOR inhibition that is independent of glucose-TOR signaling in a green alga (Fig. 8). We show that TOR inhibition blocks growth even when glucose is available, suggesting that TOR activity is required for glucose signaling. A key aspect of TOR signaling is TOR-dependent phosphorylation of target proteins, and using phosphoproteomics analysis, we report known and novel conserved targets (direct and indirect) in this pathway. Analysis of our multiomics datasets led us to postulate transcriptional and proteomic upregulation of amino acid biosynthesis pathways during TOR inhibition. Transcriptomics and proteomics also identified highly upregulated TFs. A DAP-seq experiment confirmed a bHLH TF is capable of binding to nitrogen metabolism-associated genes and thereby potentially upregulating their transcription during TOR inhibition. While previous studies showed amino acid accumulation upon TOR inhibition in photosynthetic organisms (19,21), we propose a previously unknown transcriptional regulation of nitrogen metabolism that leads to amino acid accumulation in TOR-inhibited cells. We used a *hxk1-1* mutant to distinguish TOR from the glucose-HXK1 signaling based amino acid accumulation. We conclude that regulation of nitrogen metabolism by TOR, an ancient and conserved protein, may help photosynthetic organisms maintain their carbon and nitrogen balance to support their growth and development.

**Fig. 8.**
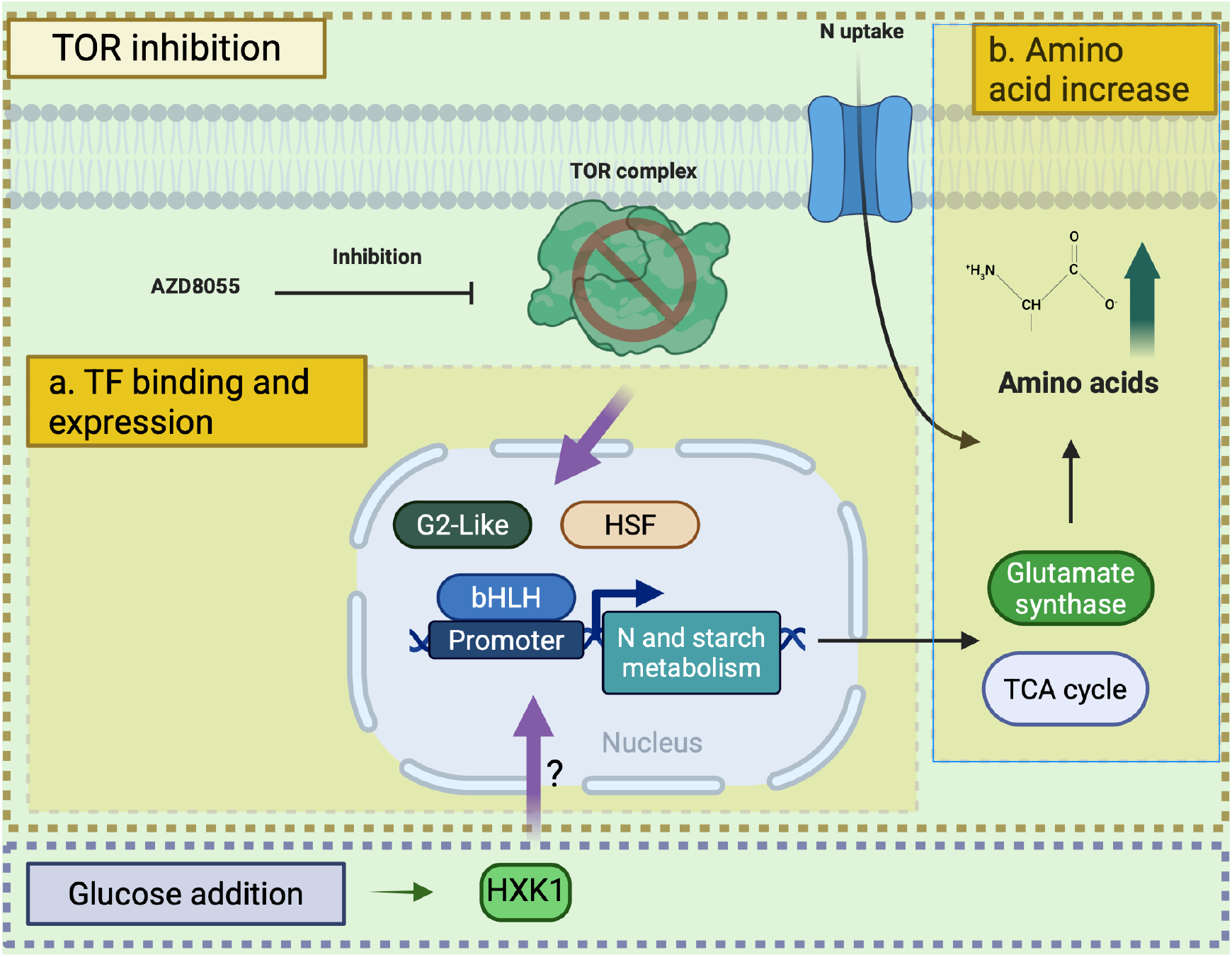
Summary of the mechanism of TOR inhibition-dependent accumulation of amino acids in *C. zofingiensis*. TOR inhibition by AZD induces the expression of transcription factors G2-like, HSF, and bHLH, which bind to amino acid biosynthesis and nitrogen metabolism genes to potentially regulate their transcription. This process, which is independent of glucose-TOR signaling, leads to an accumulation of amino acids during TOR inhibition. Glucose addition via hexokinase1 (HXK1) also upregulates amino acid biosynthesis.

### Materials and Methods

*Chromochloris zofingiensis* SAG211-14 cultures (22) were grown in *Chromochloris* Optimized Ratio of Elements CORE medium (28) under continuous 100 µmol photons m^-2^ s^-1^ light. Growth and photosynthesis metrics (cell density, F_*v*_/F_*m*_) were measured alongside omics to correlate physiological responses with molecular changes under varying nutrient and stress conditions. Samples were collected at 1 h and 24 h for multiomics analyses. Transcriptomics utilized RNA-Seq via Illumina sequencing, quantified with Salmon and DESeq2. Proteomics and phosphoproteomics involved TMT isobaric labeling, phosphopeptide enrichment, and Orbitrap mass spectrometry. Metabolomics employed LC/MS with GNPS annotation. DAP-seq identified transcription factor-DNA interactions through in vitro binding assays and motif analysis. Data integration included GO and pathway enrichment, *k*-means clustering, and orthology comparisons to identify conserved regulatory motifs. A detailed description of all methods and materials used in this study is provided in *SI Appendix*, Materials and Methods.

## Data, Materials, and Software Availability

LC-MS/MS raw data have been deposited at GNPS2https://gnps2.org/result?task=5517e1d004334a4b88f2a79a5bb0f602&viewname=filesummary&resultdisplay_type=task. RNA-Seq and DAP-seq data are available from the NCBI Gene Expression Omnibus (accession no. GSE309591). Source data are provided with this paper.

## Acknowledgments

This work was supported by the U.S. Department of Energy (DOE), Office of Science, Office of Biological and Environmental Research Award DESC0018301 (M.S.L, T.R.N., K.K.N, and M.S.R). Transcriptomics, proteomics, phosphoproteomics, and DAP-seq were performed under the Facilities Integrating Collaborations for User Science (FICUS) Program Proposals 49960 (DOI: 10.46936/fics.proj.2017.49960/ 60000021, to K.K.N and M.S.R) and used resources at the Joint Genome Institute (JGI) and the Environmental Molecular Sciences Laboratory (EMSL), which are DOE Office of Science User Facilities. Both facilities are sponsored by the Biological and Environmental Research program. Metabolomics analyses were supported by the U.S. Department of Energy, Office of Science, Office of Biological & Environmental Research, Genomic Sciences Program and used resources of the National Energy Research Scientific Computing Center, a Department of Energy Office of Science User Facility operated under contract number DE-AC02-05CH11231 to Lawrence Berkeley National Laboratory. This material was partially supported by the U.S. Department of Energy, Office of Science, Office of Biological and Environmental Research, under Award Number DE-SC0023027 (M.S.L, T.R.N., K.K.N, and M.S.R). This article is subject to HHMI’s Open Access to Publications policy. HHMI lab heads have previously granted a nonexclusive CC BY 4.0 license to the public and a sublicensable license to HHMI in their research articles. Pursuant to those licenses, the author-accepted manuscript of this article can be made freely available under a CC BY 4.0 license immediately upon publication. K.K.N. is an investigator of the Howard Hughes Medical Institute.

## Author Contributions

S.U. and M.S.R. designed research; S.U., S.M.K., A.S., R.M., Y.Z., and M.S.R. performed research; S.U., S.M.K., K.H., R.O.M., L.B., I.K.B., M.S.L., T.R.N., K.K.N., and M.S.R. analyzed data; S.U. and M.S.R. wrote the paper.

## Competing Interest Statement

The authors declare no competing interest.

## Notes

### Competing Interest Statement

The authors have declared no competing interest.

