## Supporting Figures for "TOR inhibition drives accumulation of amino acids through transcriptional activation in algae"

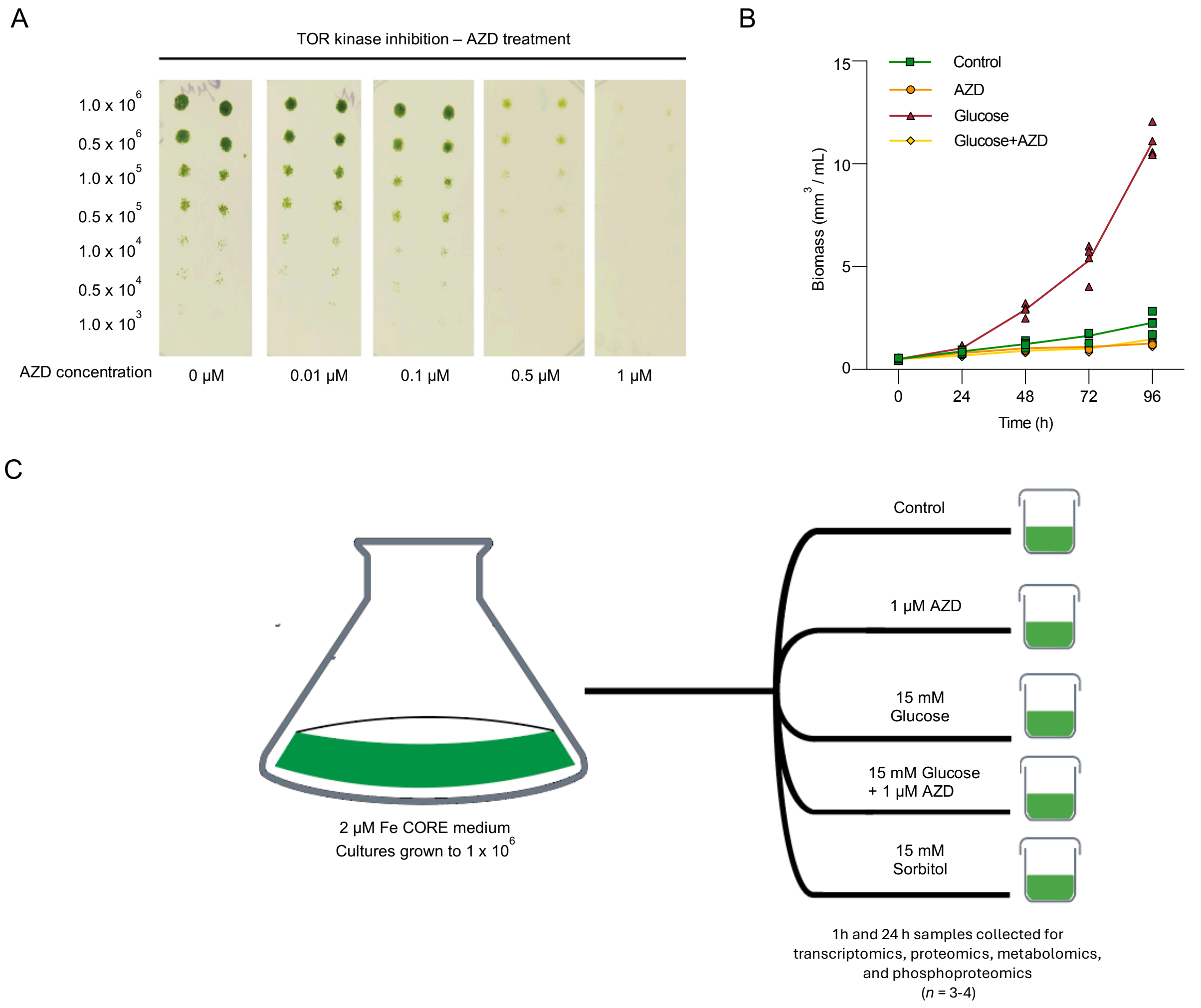


**Fig. S1.** TOR inhibition titration and the experimental outline. (A) Wild-type *C. zofingiensis* cells were subjected to serial dilutions and spotted on CORE medium plates containing the indicated concentrations of AZD8055 (AZD). Plates were incubated at 25°C under continuous illumination (100 μmol photons $m^{-2}$ $s^{-1}$). (B) Time course of volumetric culture biomass. Data represent means of *n* = 3 biological replicates. Cell density shown in Fig. 1. (C) The experimental design of multiomics experiment includes the following treatments: control, 1 μM AZD, 15 mM glucose, 15 mM glucose + 1 μM AZD, and 15 mM sorbitol (as osmotic control). Samples were collected at 1 h and 24 h.

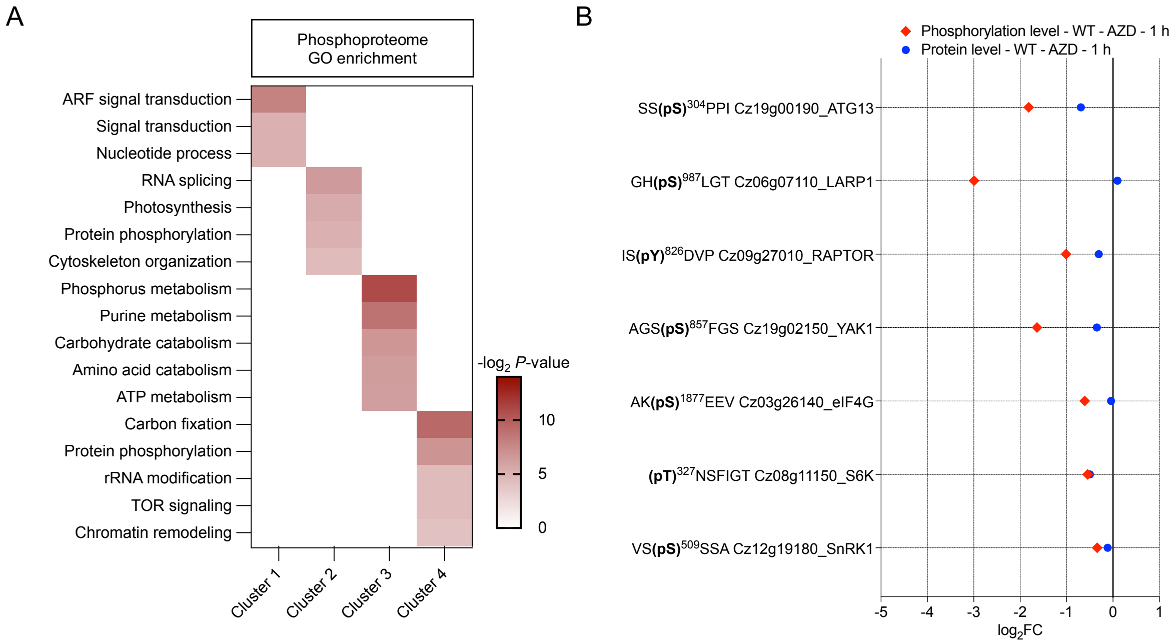


**Fig. S2.** TOR inhibited phosphoproteome enriched pathways. (A) GO term enrichment of the phosphoproteome clusters in Fig. 2A (*SI Appendix*, Dataset S2, Dataset S3). (B) $Log_2$ fold change (FC) of phosphorylated peptide and total protein for known conserved TOR-related proteins from Fig. 2B and labelled according to their protein symbols and phosphorylation Ser/Thr residue followed by their ortholog name. For (A) and (B), data represent fold changes calculated from means of $log_2$ intensities of phosphopeptides for $n$ = 4 biological replicates.

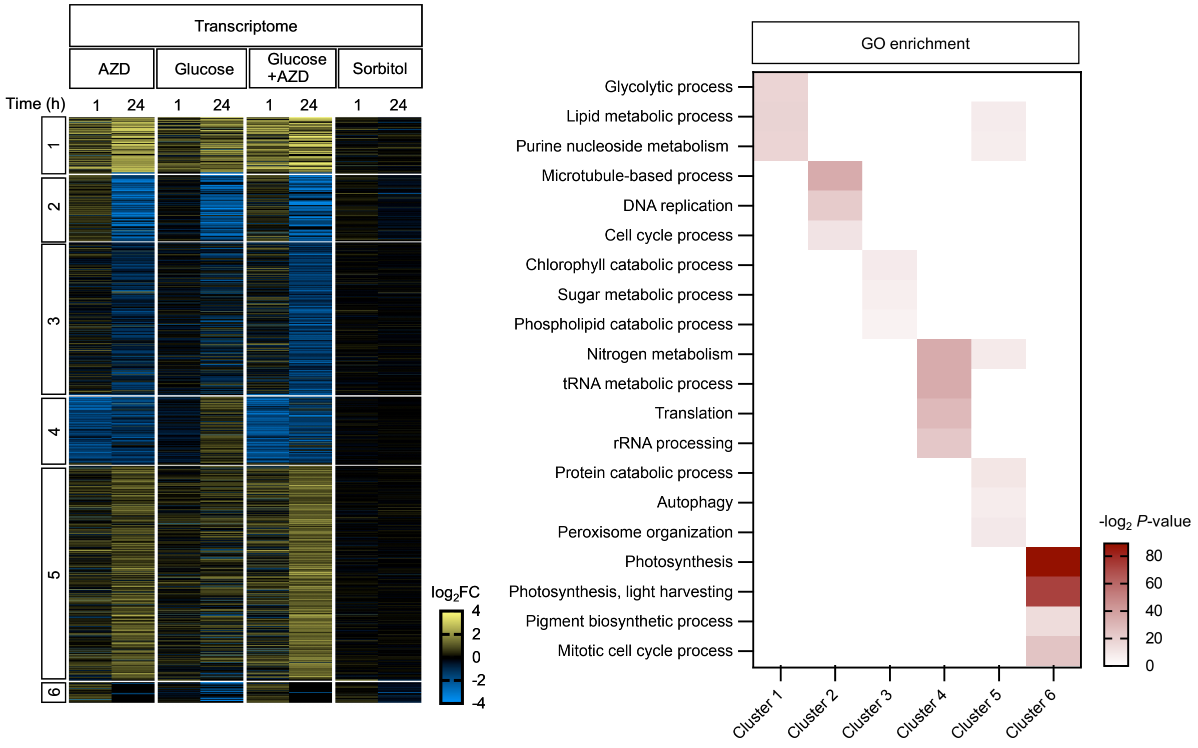


**Fig. S3.** Transcriptional regulation of global translation and protein biosynthesis genes by glucose-TOR signaling. Transcriptome $\log_2$ fold change (FC) under treatment conditions of 1 μM AZD, 15mM glucose, and 15 mM glucose+ 1 μM AZD samples was clustered using *k*-means clustering. Cluster 4 represents TOR associated genes. GO term enrichment was performed for the six transcriptome clusters. Data represent fold changes calculated from means of $\log_2$ counts of mRNA for $n$ = 3-4 biological replicates.

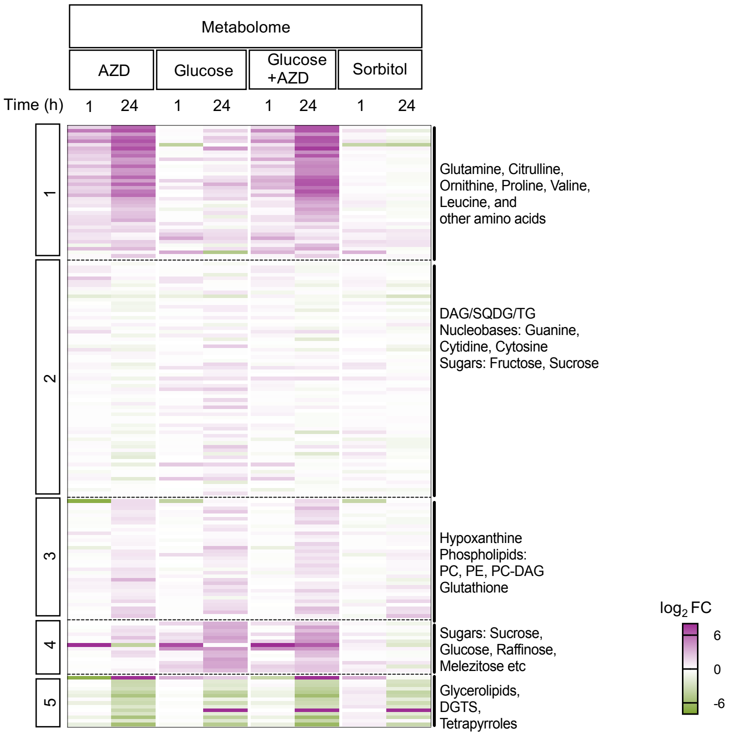


**Fig. S4.** TOR inhibition leads to increase in amino acids independent of glucose. Metabolome $\log_2$ fold change (FC) under treatment conditions of 1 μM AZD, 15 mM glucose, and 15 mM glucose + 1 μM AZD samples was clustered using *k*-means clustering. Clusters were annotated based on the metabolite classifications from GNPS (Materials and Methods). Data represent fold changes calculated from means of $\log_2$ intensities of metabolites for *n* = 4 biological replicates.

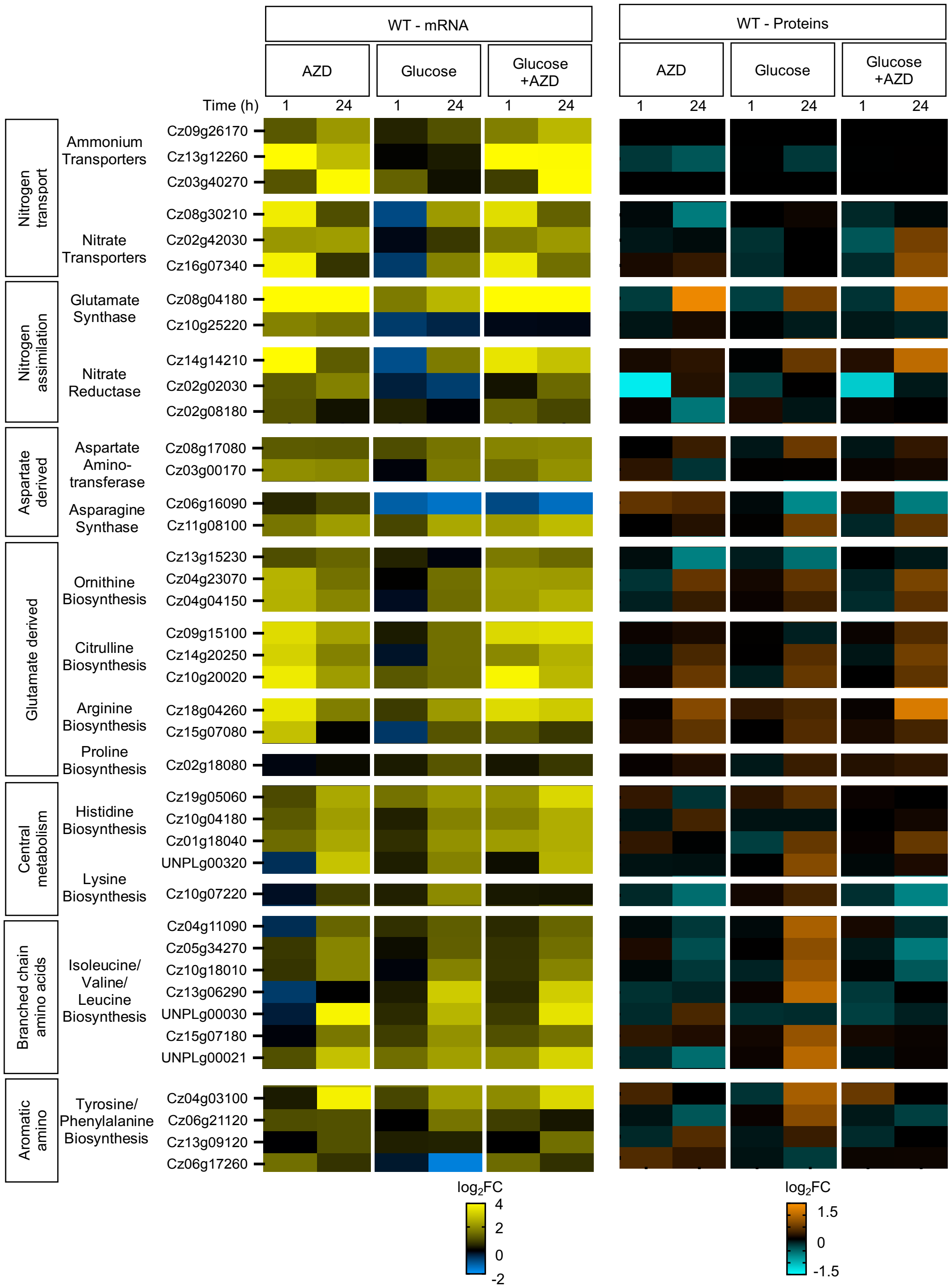


**Fig. S5.** Upregulation of de novo amino acid biosynthesis pathways during TOR inhibition in WT. Heatmap of $\log_2$ fold change (FC) of nitrogen assimilation and de novo amino acid biosynthesis genes from transcriptome and proteome during TOR inhibition by 1 μM AZD treatment, 15 mM glucose addition, and combined 15 mM glucose + 1 μM AZD treatment. The annotations for amino acid genes were obtained from previously published literature (16). Data represent fold changes calculated from means of $\log_2$ counts of mRNA and $\log_2$ intensities of proteins for $n$ = 3-4 biological replicates.

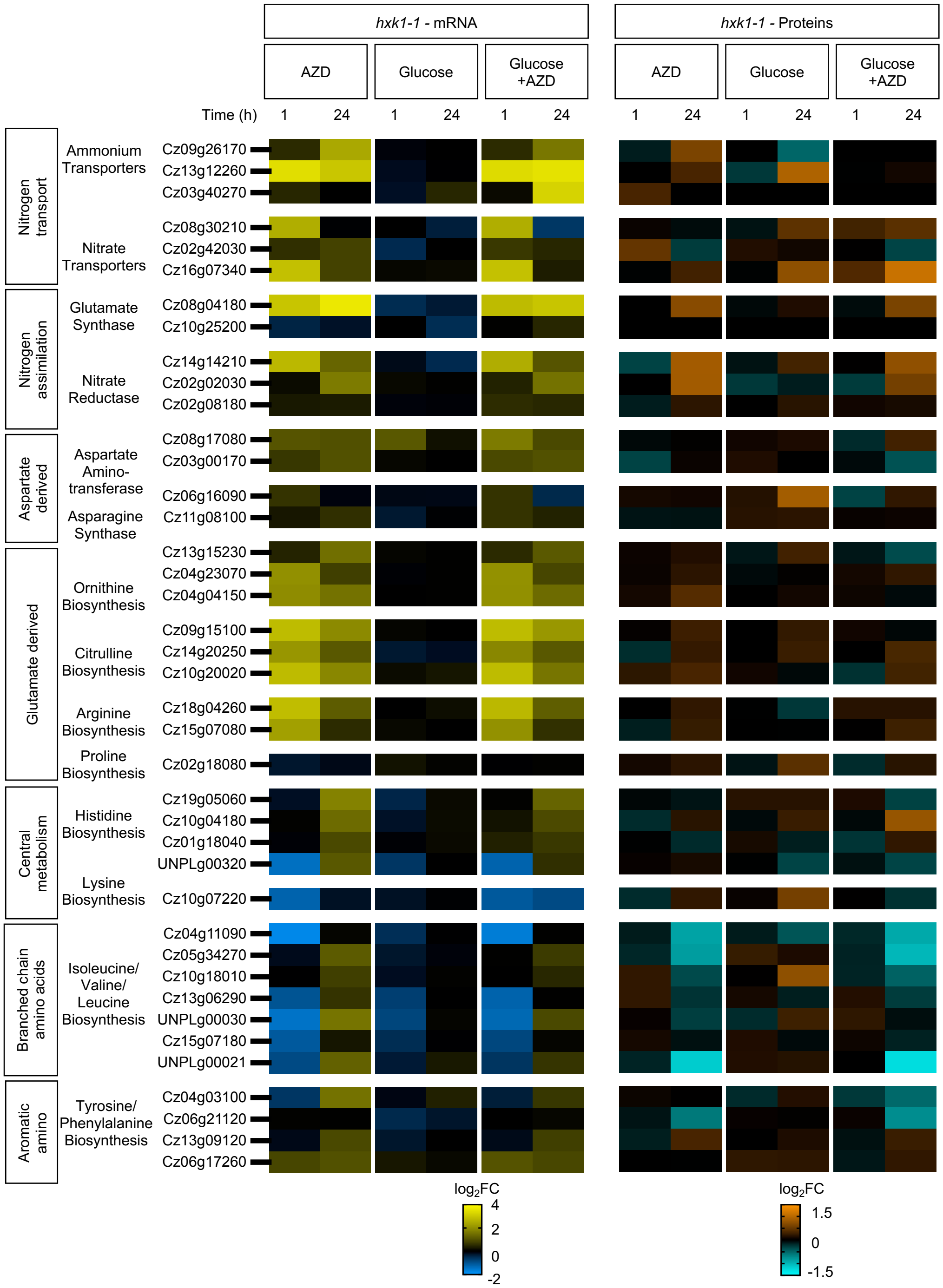


**Fig S6:** Upregulation of de novo amino acid biosynthesis pathways during TOR inhibition in *hxk1-1* mutant. Heatmap of $\log_2$ fold change (FC) of nitrogen assimilation and de novo amino acid biosynthesis genes from *hxk1-1* transcriptome and proteome during TOR inhibition by 1 µM AZD treatment, 15 mM glucose addition, and combined 15 mM glucose + 1 µM AZD treatment. The annotations for amino acid genes were obtained from previously published literature (16). Data represent fold changes calculated from means of $\log_2$ counts of mRNA and $\log_2$ intensities of proteins for $n$ = 3-4 biological replicates.

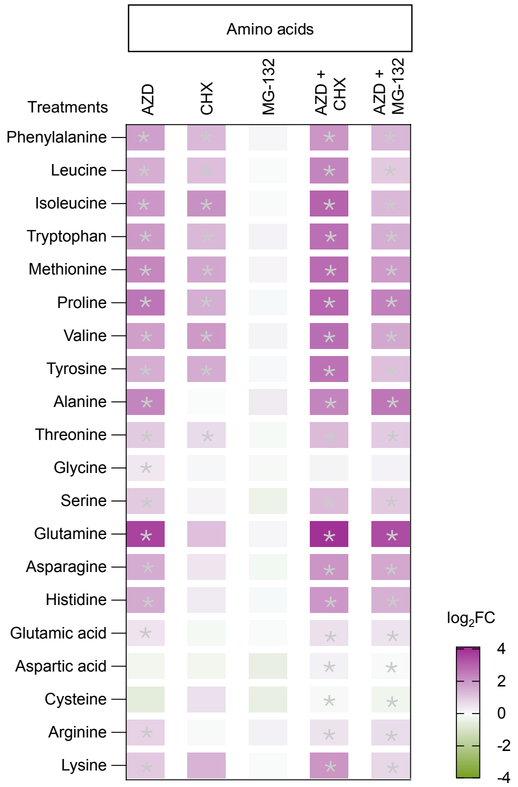


**Fig S7:** TOR inhibition leads to amino acid accumulation. Heatmap of $\log_2$ fold change (FC) of different amino acids under conditions of untreated control, TOR inhibition by 1 μM AZD, 35 μM cycloheximide (CHX), 10 μM MG-132, 1 μM AZD + 35 μM CHX, and 1 μM AZD + 10 μM MG-132 after 1 h of treatment were plotted. Data represent mean fold changes of $\log_2$ intensities of metabolites ($n$ = 3 biological replicates). Cells with asterisk represent significant differences according to one-way ANOVA and Dunnett's test: $P$<0.01.

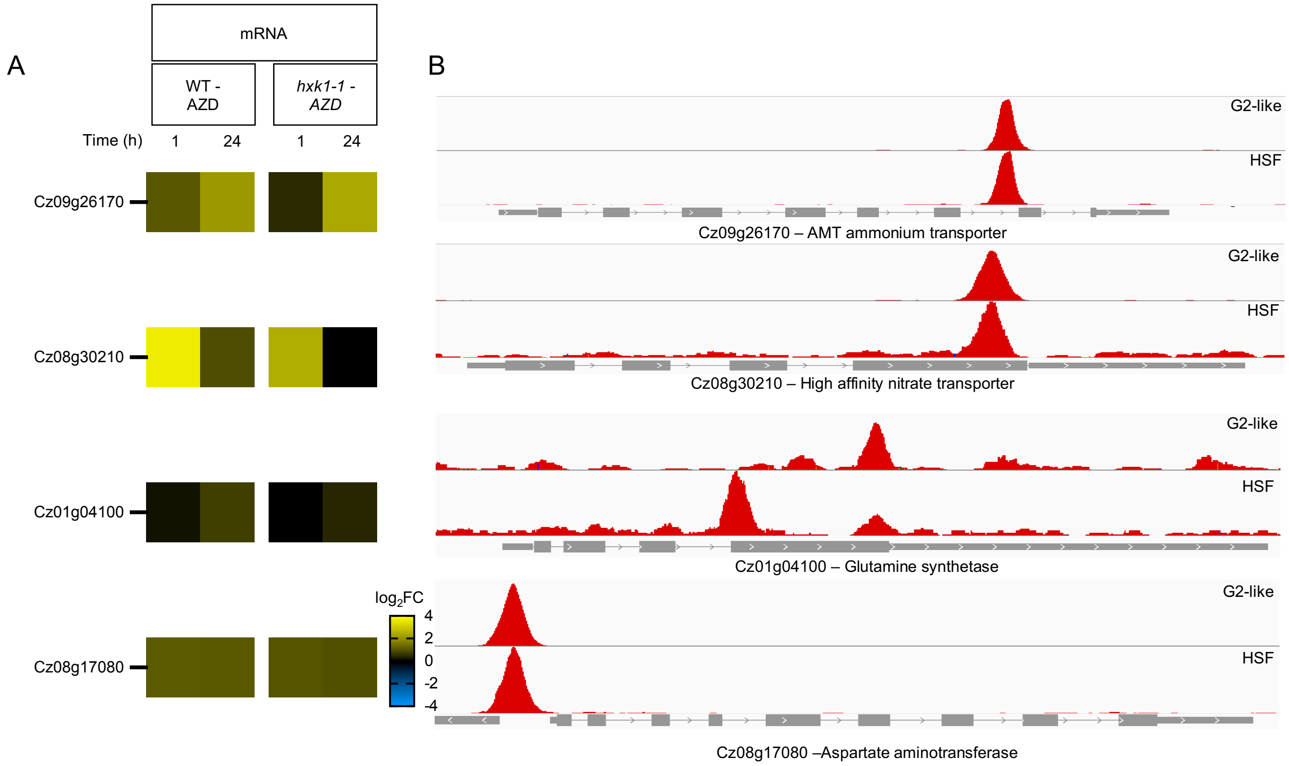


**Fig S8:** Binding of G2-like and HSF TFs to nitrogen metabolism genes potentially regulates transcription. (A) Heatmap of nitrogen metabolism genes during TOR inhibition in WT and *hxk1-1* mutant at 1 h and 24 h. Data represent fold changes (FC) calculated from means of $\log_2$ counts of mRNA for $n$ = 3-4 biological replicates. (B) DAP-seq peaks in red associated with nitrogen metabolism genes. Gene architecture with gray boxes indicates exons and lines indicate introns and gaps indicate intergenic regions.
