## Supporting Information for "TOR inhibition drives accumulation of amino acids through transcriptional activation in algae"

*Melissa S Roth

**This PDF file includes:**

Material and Methods
Figures S1 to S9
Legend for Datasets S1 to S15
References

**Other supporting materials for this manuscript include the following:**

Datasets S1 to S15

### Materials and Methods

**Strains, Growth Conditions, and sample collection for multiomics**. *Chromochloris zofingiensis* SAG211-14 cultures (1) were grown on *Chromochloris* Optimized Ratio of Elements (CORE) medium specific for optimum growth (2). For all liquid cultures, 1 x $10^5$ cells $mL^{-1}$ were inoculated in 1 L media containing 2 µM iron (Fe) in CORE media at 25$^o$C, 100 µmol photons $m^{-2}$ $s^{-1}$ and grown until 1-2 X $10^6$ cells $mL^{-1}$ density before splitting into beakers. After overnight acclimation to beakers, replicate samples were treated with either 1 µM AZD, 15 mM glucose, or 15 mM sorbitol. Sorbitol was used as a control to distinguish osmotic responses from glucose responses. Cultures were collected at 1 h or 24 h for transcriptomics, proteomics/phosphoproteomics, and metabolomics. For 96 h experiments, cultures were treated with 100mM glucose.

**Growth, biomass, and photosynthesis measurements**. Cell density and cell diameter for each treatment and time point were measured using the Multisizer 3 (Beckman

Coulter). Cultures were diluted 50-fold in an isotonic solution and the number and volume of cells per mL were counted using a 50 μM glass aperture tube. Further analysis was manually performed for each treatment and time point to calculate the volumetric biomass and cell number/mL as previously described (3).

For photosynthetic measurements, chlorophyll fluorescence of all samples was assayed using a Hansatech FMS2 system as previously described (4, 5). Samples were dark adapted for 30 min in 15 mL falcon tubes. Using a syringe filter, equal biomass of culture for each treatment were filtered on a glass fiber filter and the $F_v/F_m = (F_m - F_o)/F_m$ or maximum quantum efficiency of PSII was measured by placing the filter on a leaf clip.

**RNA preparation.** For RNA extraction, cells were collected at different time points by centrifugation (3200 g, 5 min at 4°C) and frozen in liquid nitrogen. RNA extraction and purification was performed as previously described with certain modifications (6). Briefly, 50 mL culture was pelleted via centrifugation (3,200 *g*, 5 min, 4°C) and the pellet was flash frozen in liquid nitrogen followed by fragmentation with a spatula on dry ice. Cells are then suspended in cold ethanol, transferred to Lysing Matrix D tubes, homogenized using the MP Biomedicals FastPrep-24™ 5G homogenizer (60 s, 6.5 m/s, 3 cycles), and lysed in fresh buffer containing SDS/proteinase K. The lysate is clarified by centrifugation, mixed with TRIzol (Thermo Scientific) and centrifuged to separate phases; the aqueous phase is combined with ethanol and loaded onto miRNeasy columns (Qiagen) under vacuum. After on-column DNase digestion (15 min, 30°C) RNA was eluted in water, precipitated with sodium acetate and ice-cold ethanol, incubated (−20°C, 30 min), and pelleted by centrifugation. The RNA pellet is washed with 70% ethanol, air-dried, and resuspended in RNase-free water. Finally, RNA quality is assessed via NanoDrop™ (Thermo Scientific) spectrophotometry and a 2100 Bioanalyzer (Agilent) (RIN > 8, distinct 18S/28S peaks).

**RNA quality assessment.** The protocol followed the Qiagen Prep wash and elution steps with the substitution of microcentrifuge (30–60 seconds at 14,000 rpm) in place of a vacuum manifold. RNA quality and quantity were evaluated with an Agilent 2100 Bioanalyzer. Around 3 μg RNA was submitted to Joint Genome Institute in NovoSeq S4 plates. Further steps involved creation of polyA selection-based cDNA library for 2X151 paired end read sequencing using Illumina NovoSeq. Following quality trimming of reads using the in-house JGI program BBDUK, samples had an average rRNA contamination of 0.3% and a genome mapping rate of 99.57%.

**RNA-Seq analysis**. The raw read fastq files were realigned to the updated genome annotation file published in (Cz.v.5.3.annotations.gff3, (7)). For further analysis, a decoy aware transcriptome index was generated with the reference genome v5.3 using the salmon-index generator tool. Reads were quantified from fastq files using the salmon pseudoalignment tool (v1.10.2) and the GC bias of the genome was incorporated by passing the -gcBias flag in salmon. The transcripts per million counts obtained from salmon were then used for differential expression analysis using the DESeq2 package (v1.46.0) in R. PCAs of the regularized $log_2$-transformed counts for the 500 genes with the highest variance were performed as discussed below. Heatmaps were generated in R with the ggplot2. k-means clustering was performed in R with the amap package (v0.8-14). Pathway annotations were downloaded from Phytozome's BioMart (https://phytozome.jgi.doe.gov/biomart) and functional enrichment was performed in R using the hypergeometric distribution with BH multiple testing correction and a false discovery rate of 0.05. Heatmaps were generated in Prism 10 using the $log_2$FC values obtained from DESeq2.

**Proteomics sample preparation**. Proteomics sample preparation was performed as previously described (3). Briefly, ~50 mL of culture was centrifuged at 4000 rpm for 5 min at 4$^o$C. The supernatant was discarded and the pellet was flash frozen in liquid nitrogen. Pellets were shipped in dry ice and stored at -80 °C. For protein extraction, each cell pellet was resuspended and disrupted in a bead mill homogenizer (OMNI International). After bead beating, the lysate was immediately placed on ice. To separate the proteins, metabolites, and lipids, cold chloroform:methanol (2:1) was pipetted into a microcentrifuge tube on ice and the sample was vortexed followed by centrifugation at 4°C. The protein interface had 1 mL of cold 100% methanol added to each sample, was vortexed and centrifuged again to pellet the protein. The methanol was then decanted off, and the samples were placed open in a fume hood to dry.

The protein pellet was dissolved in 8 M urea and vortexed into solution and the protein concentration was determined using a bicinchoninic acid (BCA) assay (Thermo Scientific). Following the assay, 10 mM dithiothreitol (DTT) was added, and the samples were incubated at 60°C with constant shaking at 800 rpm. Samples were then diluted 8-fold in preparation for trypsin digestion. 100 mM $NH_4HCO_3$, 1 mM $CaCl_2$ and sequencing-grade modified porcine trypsin (Promega) were added to all protein samples at a 1:50 (w/w) trypsin-to-protein ratio for 3 h at 37℃ with constant shaking at 450 rpm. Digested samples were desalted using a 4-probe positive pressure Gilson GX-274 ASPECTM system (Gilson Inc.) with Discovery C18 100 mg/1 mL solid phase extraction tubes (Supelco), using the following protocol: 3 mL of methanol was added for conditioning followed by 2 mL of 0.1% trifluoroacetic acid (TFA) in $H_2O$. The samples were then loaded onto each column followed by 4 mL of 95:5: $H_2O$: acetonitrile, 0.1% TFA. Samples were eluted with 1 mL 80:20 acetonitrile:$H_2O$, 0.1% TFA. The samples were concentrated down to ~100µL using a SpeedVac, and a final BCA assay was performed to determine the peptide concentration. An equal mass of each sample was aliquoted into fresh centrifuge tubes, dried completely in a SpeedVac and then stored at -80°C until isobaric labeling.

**TMT isobaric tag labeling.** Each sample was diluted in 500 mM HEPES, pH 8.5 to a concentration of 5 µg/µl and labeled using amine-reactive Thermo Scientific Tandem Mass Tag (TMT10) Isobaric Mass Tagging Kits (Thermo Scientific) according to the manufacturer's instructions. Briefly, 250 µL of anhydrous acetonitrile was added to each 5 mg reagent, vortexed and allowed to dissolve for 5 min with occasional vortexing. Reagents were then added to the samples and incubated for 1 h at RT with shaking at 400 rpm. Each sample was then diluted to 2.5 µg/µl with 20% acetonitrile, and the reaction was quenched by adding 8 µL of 5% hydroxylamine to the sample with incubation for 15 min at RT with shaking at 400 rpm. The samples within each set were then combined and completely dried in the SpeedVac. Each of the samples were then cleaned using C18 50 mg/1 mL solid phase extraction tubes as described above and once again assayed with BCA to determine the final peptide concentration.

**Phosphopeptide Enrichment.** The TMT labeled peptides were subjected to phosphopeptide enrichment using magnetic nickel-nitriloacetic acid (Ni-NTA) beads (Qiagen) according to the manufacturer's protocol. Ni-NTA magnetic beads were washed three times with water and treated with 100 mm EDTA, pH 8.0 for 30 min. After removing the EDTA solution, beads were washed three times with water, and treated with 10 mm $FeCl_3$ for 30 min. After removing excess metal ions, the beads were washed three times with water and resuspended in 1:1:1 (v/v/v) acetonitrile/methanol/0.01% (v/v) acetic acid. Beads were then conditioned using resuspending/wash buffer containing 80% (v/v) acetonitrile and 0.1% (v/v) trifluoroacetic acid, followed by incubation with peptides (100 µg peptides in 200 µl of 80% (v/v) acetonitrile with 0.1% (v/v) trifluoroacetic acid) for 30 min. The beads were washed three times with the resuspending/wash buffer and the phosphopeptides were eluted with elution buffer containing 50% (v/v) acetonitrile and 2.5% (v/v) ammonia. The eluate containing phosphopeptides was immediately acidified with 10% (v/v) trifluoroacetic acid and concentrated by vacuum centrifugation prior to mass spectrometry analysis.

**Offline fractionation of peptides and preparation of proteome samples.** Labeled peptides were separated using an offline high pH (pH 10) reversed-phase (RP) separation on a Waters XBridge C18 column (250 mm × 4.6 mm, 5 µm particles) with a 4.6 mm × 20 mm guard column, utilizing an Agilent 1200 HPLC System. The sample was loaded onto the column and washed for 15 minutes with Solvent A (10 mM ammonium formate, pH 10). The LC gradient began with a linear increase of Solvent B (10 mM ammonium formate, pH 10, 90:10 acetonitrile:water ) to 5% over 10 minutes, followed by an increase to 45% over 65 minutes, and then a linear increase to 100% over 15 minutes. Solvent B was held at 100% for 10 minutes before returning to 100% Solvent A for 20 minutes to recondition the column, all at a flow rate of 0.5 mL/min. A total of 96 fractions were collected into a 96-well plate, which were then concatenated into 24 fractions using a previously reported strategy (8), excluding CHAPS-containing wells. The peptide fractions were dried down, re-suspended in nanopure water at a concentration of 0.075 µg/µL, and analyzed using a Q Exactive Hybrid Quadrupole Orbitrap Mass Spectrometer (Thermo Scientific).

**Mass-Spectrometry Based Proteomic Analysis of Samples.** All peptide samples were analyzed using an automated constant-flow nano LC system (Agilent) coupled to a Q Exactive Orbitrap mass spectrometer (Thermo Fisher Scientific). Custom-made electrospray emitters were fabricated from chemically etched fused silica with dimensions of 150 µm outer diameter (OD) x 20 µm OD x 20 µm inner diameter (ID). Separation was performed on a 4-cm fused-silica capillary analytical column (360 µm OD x 150 µm ID) packed with 3 µm Jupiter C18 particles. The mobile phases consisted of 0.1% formic acid in water (solvent A) and 0.1% formic acid in acetonitrile (solvent B), delivered at a flow rate of 300 nL/min. The gradient elution profile was programmed as follows (time in minutes: %B): 0:5, 2:8, 20:12, 75:35, 97:60, and 100:85.

Mass spectrometric analysis was performed on an LTQ Orbitrap Velos (Thermo Fisher) operating in data-dependent acquisition mode. Full MS scans were acquired at a resolution of 30,000 with a target of 3 x $10^6$ ions across a mass range of 300–1800 m/z. The top ten most abundant precursor ions were selected for higher-energy collisional dissociation (HCD) with a resolution of 7,500 and a target of 5 x $10^4$ ions. An isolation window of 2.5 Th was applied prior to HCD fragmentation. HCD scans were conducted using a normalized collision energy of 45 and a maximum injection time of 1000 ms. Dynamic exclusion was set to 60 s to prevent repeated analysis of the same ions, and charge state screening was enabled to exclude unassigned and singly charged ions.

**Peptide Identification and Quantification.** Peptide identification was performed using MS/MS spectra were searched against a decoy *A. thaliana* protein database (ftp://ftp.ncbi.nlm.nih.gov/genomes/all/GCF/000/715/135/ GCF_000715135.1_Ntab-TN90/) as well as a contaminants database containing human keratin and trypsin sequences, using the algorithm SEQUEST (9). Protein fasta databases used at PNNL were derived from the *C. zofingiensis* genome (1, 7). Search parameters included: no enzyme specificity for proteome data and trypsin enzyme specificity with a maximum of two missed cleaves, ± 50 ppm precursor mass tolerance, ± 0.05 Da product mass tolerance, and carbamidomethylation of cysteines and TMT labeling of lysines and peptide N-termini as fixed modifications. Allowed variable modifications were oxidation of methionine. For phosphopeptide analysis, a variable modification of phosphorylation of serine (S), threonine (T) or tyrosine (Y) were additionally searched and the software ASCORE (10) was used to determine phosphorylation site. Measured mass accuracy and MSGF spectra probability were used to filter identified peptides to <0.4% false discovery rate (FDR) at spectrum level and <1% FDR or <5% FDR at the peptide level using the decoy approach (11). TMT reporter ions were extracted using the MASIC software (12) with a 20 ppm mass tolerance for each expected TMT reporter ion as determined from each MS/MS spectrum.

**Downstream analysis with peptide and protein abundances.** For phosphoproteome analysis, raw data files with 1% FDR were used. Contaminant rows and blank columns were removed for analysis. In the next step, rows with missing values were also removed. The resultant peptide list for control, sorbitol, glucose, AZD, and glucose+AZD were tested for statistical analysis of global differential peptide expression. For calculation of $\log_2$fold change ($\log_2$FC), the raw peptide data was converted to $\log_2$ peptide and the biological replicates were averaged followed by parametric t-tests and *p* values were also extracted using R in RStudio. The following contrasts were tested for both 1 h and 24 h: Sorbitol 1 h vs Control 1 h, AZD 1h vs control 1 h, glucose 1h vs Control 1 h, glucose+AZD 1 h vs Control 1 h, Sorbitol 24 h vs Control 24 h, AZD 24 h vs Control 24 h, Glucose 24 h vs Control 24 h, and Glucose+AZD 24 h vs Control 24 h. $\text{Log}_2$FC lists with p values were generated for each treatment and provided as supplementary data. The peptides were further filtered based on $\log_2\text{FC} \geq 1$ and $P \leq 0.01$ and used for clustering and GO enrichment analysis as described below. To identify conserved phosphopeptides, known TOR kinase target proteins that were reported previously in *C. reinhardtii* and *A. thaliana*

(13–15) were extracted using a selective literature screen and blasted against the *C. zofingiensis* genome. The identified orthologs from *C. zofingiensis* were then aligned to both *C. reinhardtii* and *A. thaliana* proteins and the conserved phosphopeptides were reported. To report conservation of phosphosites in *C. zofingiensis*, *C. reinhardtii*, *Auxenochlorella sp.* UTEX250-A and *A. thaliana*, a sequence window of 8-12 residues surrounding the phosphosite was reported for selected targets of TOR kinase.

For global proteome analysis, peptide rollup was performed for all proteins to combine the intensities of all peptides of a protein. Contaminant rows and blank columns were removed for analysis. Rows with missing values were removed and further statistical analysis was performed on the resultant table of combined intensities. For PCA analysis of the global proteome, the raw intensity values were log transformed, and percent variance was calculated using the prcomp function in R using the PCAtools package (v2.18.0). The first two principal component coordinates were then extracted and plotted through ggplot2 (v3.3.5) to provide shape and color features consistent with the variables. For statistical analysis of the global proteome, $\log_2$FC of the protein intensities were calculated for the above-mentioned contrasts and t-tests were performed in R to calculate the p value. Pearson correlation was calculated between the $\log_2$FC of transcript and protein for each gene. The correlation was plotted using Prism 10. Pathway enrichment was performed as described below.

**Metabolomic sample preparation.** For metabolite extraction, cells were pelleted at required time points for 5 min at 4000 rpm in a GeneMate 2 mL screw cap tubes. The supernatant was discarded and the cells were frozen in liquid nitrogen. Cell pellets were resuspended in 500uL water and pipetted up and down to break apart cell pellet; the suspended pellets were then immediately frozen at -80C and lyophilized. The dried material was homogenized for 5 seconds, 3 times with 10 second breaks to prevent overheating with a 3.2mm stainless steel bead (BioSpec Products) in each tube using a Mini-Beadbeater (BioSpec Products). Homogenized dried cell material was then resuspended in 150uL of methanol with internal standards (Dataset S17), vortexed 2x 10 seconds, ice bath sonicated for 15 minutes, centrifuged (10,000 rcf, 5 min, 10C), and then supernatants were filtered (0.22um pvdf) and transferred to amber glass vials for LCMS analysis. Briefly, metabolites were separated using hydrophilic liquid interaction chromatography on an Agilent 1290 HPLC and detected on a Thermo QExactive Hybrid Quadrupole-Orbitrap Mass spectrometer; LCMS parameters are available in SI Dataset S17.

**Metabolomics data analysis.** Targeted annotations of amino acids were made using Metatlas (26, 27, 28) to extract peak heights using known m/z, retention time and fragmentation spectra based on reference standards analyzed on the same method, identification were determined per the Metabolomics Standards Initiative (18); outputs including evidence of identifications and peak heights are available in Dataset S18)

Feature-based molecular networking (FBMN) was performed as described before using MZmine2 and GNPS (16, 17). Positive mode top GNPS annotations were accepted with cosine scores of MS/ MS spectral mirror match (MQScore) to library reference of >0.7. Additional filtering included removing known contaminants and adducts such as

Phthalic anhydride, Diocytl phthalate and Benzyltetradecyldimethylammonium etc. Untargeted metabolites with retention time between 1 and 17.5 and with an MZErrorPPM values of less than 10 were used. Next, the NPClassifier determined metabolite classifications based on annotated features of biosynthetic pathways. For filtered features, peak heights with differential abundances between different treatments were compared to calculate log fold change values. The $log_2FC$ values were then clustered using *k* means clustering as described below and the metabolite NPclassifier assignments from GNPS were labelled on the heatmap (Prism 10).

**Gene Ontology (GO), Pathway enrichment, and clustering analysis.** GO annotation and pathway annotations per protein were downloaded from Phytozome's BioMart. GO term enrichment was performed with the topGO package (v1) using the Fischer's exact test. Pathway enrichment was calculated in R using the enricher function that performs a hypergeometric test with BH multiple testing correction (ClusterProfiler package in R) (25). GO terms and pathways were filtered using an FDR of <= 0.05 and the lists were generated in R. *k*-means clustering was performed in R with the amap package (v0.8-14). Functional annotations for pathway analysis in Figures 4 and 5 were adapted from (19).

**DAP-seq experiment and analysis**. The DAP-seq experiment was performed as per standardized protocol at the Joint Genome Institute (20, 21). Briefly, genomic DNA (gDNA) was extracted from *C. zofingiensis* cells using a CTAB protocol, fragmented to an average size of 150 bp and ligated with custom adapters. Coding sequence (CDS) of selected TFs were cloned into the pIX-HALO-PaqCI vector directly downstream of the HaloTagHalo-tagged. The T7 promoter driven HaloTag-fused TF CDSs were amplified by using primers pIX-Halo-T7-fwd (5′-GTGAATTGTAATACGACTCACTATAGGG) and pIX-Halo-AfterPolyA-rev (5′-CAAGGGGTTATGCTAGTTATTGCTC), the resulting PCR products were purified, and the correct amplicon sizes were verified on a Fragment Analyzer system (Agilent). The TF proteins were then in vitro expressed using the TnT T7 Quick for PCR DNA system (Promega L5540) and bound to the Magne HaloTag Beads (Promega G7282) on a 96 well plate. The protein-bound beads were then incubated with *C. zofingiensis* gDNA libraries and then gently washed to remove any unbound DNA. After the final wash, the beads were resuspended in diluted index primer mix and boiled at 95°C for 10 min to elute DNA off from beads. The eluate from each well was transferred to a new plate and mixed with the KAPA HiFi HotStart ReadyMix (Roche) for amplification. The PCR products were then gel purified and sequenced using Illumina NovaSeq 2X150 bp paired-end sequencing.

The sequencing fastq files were quality filtered and adapters were trimmed using BBTools and then aligned to the reference genome of *C. zofingiensis* v5.3.2 using bowtie2 v2.4.2. The bam files were created using samtools v1.15.1 (22). The MACS3 v3.0.0a6 callpeak command was used to generate narrowPeak files, using a background control obtained from DAP-seq wells with a mock TF protein expression. Motifs were called using MEME suite v5.3.0 (23) command with the reference genome nucleotide frequency as the background model. The first positional weight matrix motif identified by MEME motif caller (MEME-1) from the summit regions of the top peaks was used. TFs that performed poorly in the DAP assay were removed. Successful binding was calculated using the fraction of reads in peaks (FRIP) scores and only TFs with FRIP scores $\geq 0.05$ were used for further analysis. The DAP-seq peaks identified in the analysis above were further processed to identify target genes for each TF. Using the gene annotation file (gff format), features corresponding to transcript/mRNA were filtered and assigned up to two gene targets. Further detailed assignment of peaks to either intron, exon, intron-exon-junction, promoter (+- 2000 bp) was done manually to plot the peak distribution for each TF. Finally, pathway enrichment was performed for the target genes of each TF using the ClusterProfiler package in R as previously described (25).

**Orthology analysis and identification of conserved motifs in promoters.** Orthologues of proteins in Fig. 6 were identified by BLAST in Phytozome for the following species: *C. reinhardtii*, *V. carteri*, *A.* sp. UTEX 250-A, *A. thaliana*, *Oryza sativa*. For *A.* sp. UTEX 250-A, access to the genome sequence and annotations was kindly provided by the Merchant group (24). To extract the upstream and downstream promoter regions, ±1500 bp from the transcription start site, geneIDs were provided in BioMart for each species and then searched for the presence or absence of the CACGTG/CANNTG motifs.

**Legend for supporting datasets:**

Dataset S1 (separate file). DatasetS1.xlsx. Detection of 19,030 phosphopeptides on 3861 proteins with a peptide being detected in at least one treatment.

Dataset S2 (separate file). DatasetS2.xlsx. Clustering of $log_2$FC of 1288 significant phosphopeptides at different treatment conditions of AZD, glucose, glucose+AZD, and sorbitol.

Dataset S3 (separate file). DatasetS3.xlsx. GO term enrichment of the phosphopeptide clusters in Fig. S2A.

Dataset S4 (separate file). DatasetS4.xlsx. Clustering of $log_2$FC of transcripts under different treatment conditions.

Dataset S5 (separate file). DatasetS5.xlsx. GO term enrichment of 6 clusters identified in the transcriptome.

Dataset S6 (separate file). DatasetS6.xlsx. Comparison of the $log_2$FC of 9528 genes between the transcriptome and proteome datasets of AZD, glucose, and glucose+AZD treated samples at 24 h.

Dataset S7 (separate file). DatasetS7.xlsx. Pathway enrichment of sectors identified in Fig. 3.

Dataset S8 (separate file). DatasetS8.xlsx. Clustering of $log_2$FC of WT metabolites under different treatments.

Dataset S9 (separate file). DatasetS9.xlsx. $Log_2$FC of WT transcripts and proteins associated with nitrogen metabolism and de novo amino acid biosynthesis pathways.

Dataset S10 (separate file). DatasetS10.xlsx. $Log_2$FC of *hxk1-1* mutant transcripts and proteins associated with nitrogen metabolism and de novo amino acid biosynthesis pathways.

Dataset S11 (separate file). DatasetS11.xlsx. $Log_2$FC of metabolites after 1 h of treatment with no treatment (control), AZD, cycloheximide, MG-132, AZD+cycloheximide, and AZD+MG-132.

Dataset S12 (separate file). DatasetS12.xlsx. $Log_2$FC of differentially expressed TFs from clusters 1 and 5 of the transcriptome (Fig. S3).

Dataset S13 (separate file). DatasetS13.xlsx. Target genes of three TFs.

Dataset S14 (separate file). DatasetS14.xlsx. Pathway enrichment of target genes associated with the three TFs.

Dataset S15 (separate file). DatasetS15.xlsx. Orthologues of target genes related to nitrogen and starch metabolism with the E/G-box identified in their promoters.

Dataset S16 (separate file). DatasetS16.xlsx. GNPS output features for filtered metabolites.

Dataset S17 (separate file). DatasetS17.xlsx. LC MS Parameters.

Dataset S18 (separate file). DatasetS18.xlsx. Targeted annotations of amino acids.
